# Multilayered mechanisms ensure that short chromosomes recombine in meiosis

**DOI:** 10.1101/406892

**Authors:** Hajime Murakami, Isabel Lam, Jacquelyn Song, Megan van Overbeek, Scott Keeney

## Abstract

To segregate accurately during meiosis, homologous chromosomes in most species must recombine. Very small chromosomes would risk missegregation if recombination were randomly distributed, so the double-strand breaks (DSBs) that initiate recombination are not haphazard. How this nonrandomness is controlled is not understood. Here we demonstrate that *Saccharomyces cerevisiae* integrates multiple, temporally distinct pathways to regulate chromosomal binding of pro-DSB factors Rec114 and Mer2, thereby controlling duration of a DSB-competent state. Homologous chromosome engagement regulates Rec114/Mer2 dissociation late in prophase, whereas replication timing and proximity to centromeres or telomeres influence timing and amount of Rec114/Mer2 accumulation early. A distinct early mechanism boosts Rec114/Mer2 binding quickly to high levels specifically on the shortest chromosomes, dependent on chromosome axis proteins and subject to selection pressure to maintain hyperrecombinogenic properties of these chromosomes. Thus, an organism’s karyotype and its attendant risk of meiotic missegregation influence the shape and evolution of its recombination landscape.

## Introduction

A critical goal in meiotic prophase is to allocate crossovers to all pairs of homologous chromosomes to ensure proper segregation in the first meiotic division (MI) (Page and Hawley, 2003). Crossovers are produced by homologous recombination initiated by programmed DNA double-strand breaks (DSBs). However, DSBs could also have deleterious consequences, so their formation is controlled by multiple regulatory pathways to ensure that they occur at the appropriate time, number, and place to avoid genome instability (Keeney et al., 2014). The full array of these pathways is not yet known, nor is it understood how they integrate with one another to promote essential functions of DSBs while minimizing the risk.

In budding yeast, DSB formation requires topoisomerase-like Spo11 and nine accessory proteins (“DSB proteins,” including Rec114 and Mer2), which interact with each other and assemble complexes on meiotic chromosomes (Arora et al., 2004; Li et al., 2006; Maleki et al., 2007; Panizza et al., 2011). Most of these factors are nearly universal in eukaryotes, including fission yeast (Miyoshi et al., 2012) and mammals (Kumar et al., 2010; Robert et al., 2016; Stanzione et al., 2016; Tesse et al., 2017).

The timing of DSB formation is regulated at multiple levels. For example, six of ten DSB proteins are only expressed in meiosis (Keeney, 2007). Furthermore, recruitment of Rec114 depends on phosphorylation of Mer2 by cell cycle regulatory kinases CDK and DDK (Henderson et al., 2006; Murakami and Keeney, 2008; Sasanuma et al., 2008; Wan et al., 2008), and DDK phosphorylation is directly coupled with replication fork passage (Murakami and Keeney, 2014).

The control of DSB number per chromosome is also vital, because homologous chromosomes can only recombine if they have at least one DSB between them. To illustrate this point, we performed a Monte Carlo simulation of DSB formation in *S. cerevisiae,* which makes an esti-mated 150–200 DSBs per meiosis (Pan et al., 2011). We randomly distributed 200 DSBs across the 16 yeast chromosomes in proportion to their length and assessed how often a cell failed to get at least one DSB on every chromosome. From 10,000 simulations, 398 (4%) had at least one chromosome that failed to form a DSB. Not surprisingly, the three shortest chromosomes (chr1, 3 and 6) were by far the most prone to failure, accounting for 98% of total mishaps (**Figure 1A**).

**Figure 1.**
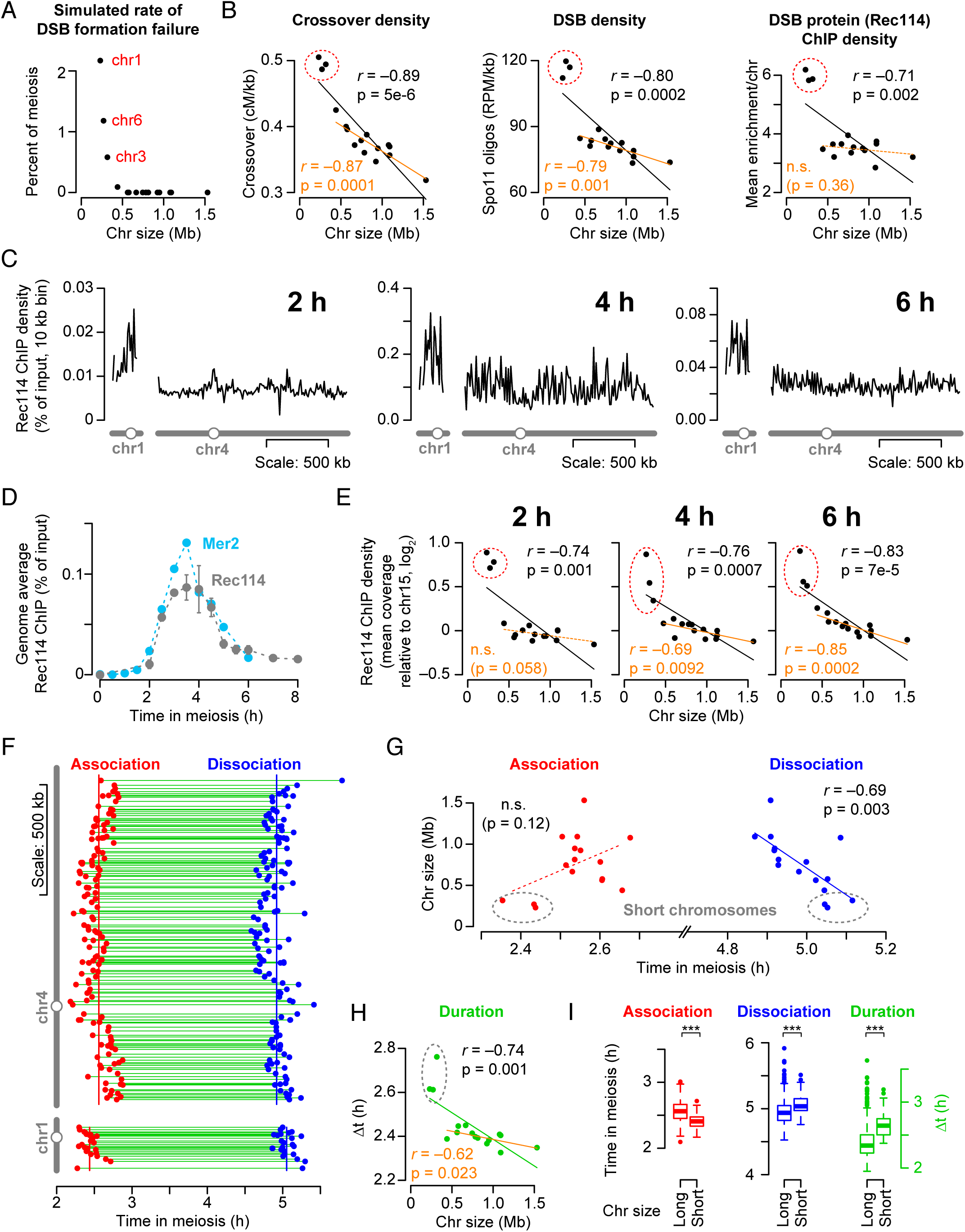
Rec114 and Mer2 accumulate preferentially on smaller chromosomes. (A) Simulation of randomly distributed DSBs. For each simulated meiosis, 200 DSBs were distributed randomly according to chromosome size. The y axis shows the percent of simulations (10,000 total) in which the indicated chromosome failed to acquire a DSB. (B) Chromosome size dependence of crossovers (centiMorgans (cM) per kb), DSBs (Spo11-oligo density in reads per million (RPM) per kb), and Rec114 binding (ChIP-chip enrichment). In all figures, chr12 is represented without including its full rDNA cluster. Data are from (Mancera et al., 2008; Pan et al., 2011; Panizza et al., 2011). (C) Example Rec114 ChIP-seq profiles for chr1 and chr4 at the indicated times. Absolute ChIP signal was calibrated by qPCR and smoothed with 10-kb sliding window. (D) Time course of genome-average Rec114 and Mer2 levels. Gray points are mean ± range of the two Rec114 datasets. Cyan points are from the single Mer2 dataset. (E) Size dependence of average per-chromosome Rec114 ChIP density. For each time point, the mean ChIP signal for each chromosome was normalized to the mean for chr15 and log2-transformed. The full time course is in Figure S1B. (F) Within-chromosome organization of Rec114 association and dissociation times. Each point is a called Rec114 peak. Green lines indicate binding duration (δt). (G) Per-chromosome mean association and dissociation times for Rec114. (H) Per-chromosome mean Rec114 binding duration. (I) Rec114 shows early association and late dissociation on short chromosomes and thus longer duration of binding. Values for Rec114 peaks were grouped for the shortest three chromosomes (n= 71 peaks) and the other thirteen chromosomes (“long”, n= 927 peaks). In all boxplots: thick horizontal bars are medians, box edges are upper and lower quartiles, whiskers indicate values within fold of interquartile range, and points are outliers. *** indicates p < 0.005, one-sided Wilcoxon test. Panels C and E–I are from the *ARS+* time course; other time courses are in Figure S1. See also Figure S1.

This simple exercise indicates that the shortest chromosomes would be at especially high risk of meiosis I nondisjunction if DSBs were randomly distributed, but several lines of evidence indicate that there are mechanisms that attenuate this risk. Specifically, it has long been known that crossover density (centimorgans per kb) negatively correlates with chromosome size (Kaback et al., 1992; Chen et al., 2008; Mancera et al., 2008) (**Figure 1B**). This correlation may be related to chromosome size per se because crossover density can sometimes but not always be altered by changing chromosome length (Kaback et al., 1992; Turney et al., 2004). A negative correlation with chromosome size is also observed for DSB density (Pan et al., 2011; Thacker et al., 2014) and for the amount of DSB protein binding assessed by chromatin immunoprecipitation (ChIP) (Panizza et al., 2011; Sun et al., 2015) (**Figure 1B**). These observations suggest that short chromosomes recruit more DSB proteins compared to long chromosomes, leading to a higher density of DSBs and thus crossovers. How this preferential DSB protein recruitment is achieved is largely unknown.

In this study, we show how multiple mechanisms play temporally distinct roles to regulate the time of chromosomal association and dissociation of DSB proteins, thereby controlling the duration of a presumptive DSB-competent state. We also show that a separable mechanism allows the shortest chromosomes to quickly accumulate especially high levels of DSB proteins, and we provide evidence that selective pressure maintains this property of little chromosomes over evolutionary time scales. Our results create an integrated view of a multilayered and evolutionarily constrained system that ensures allocation of DSBs and thus crossovers to all pairs of homologous chromosomes.

## Results

### Rec114 and Mer2 are present at higher levels and for longer times on short chromosomes

To understand how DSB proteins associate with meiotic chromatin, we performed ChIP-seq for Rec114 or Mer2 at multiple time points from three synchronized meiotic cultures and calibrated coverage maps by quantitative PCR (qPCR (Murakami and Keeney, 2014)) (**Figures 1C and 1D**). Two Rec114 datasets were described previously (Murakami and Keeney, 2014). In one, major replication origins on the left arm of chr3 (chr3L) were deleted (*arsΔ*) to determine the effect of replication delay; in the other, origins were intact. For this study, we added a third time course assessing Mer2 binding in an *arsΔ* strain. Rec114 and Mer2 coverage maps agreed well throughout meiosis (**Figure S1A**), suggesting they are regulated similarly. The effect of replication delay was also similar (see below).

ChIP signal was essentially zero at the beginning of the time courses as expected, was low but readily detectable at 2 h (during replication), peaked at ∼3 to 4 h (coinciding with peak steady-state DSB levels), and decreased thereafter (**Figures 1C and 1D**). Because absolute ChIP signal varied widely over time, we used the second longest chromosome (chr15) as an internal normalization control to evaluate relative per-chromosome abundance (**Figures 1E, S1B, and S1C**). Two key patterns emerged. First, ChIP signal was overrepresented on short chromosomes throughout meiotic prophase. Second, early and late per-chromosome ChIP densities showed markedly different patterns. At early time points (2–3 h) the three shortest chromosomes stood out starkly as having high relative ChIP densities, while the rest had low ChIP densities with no size correlation. Later (4.5–6 h), the 13 largest chromosomes showed a clear negative correlation with size and the smallest three deviated less from an overall linear relationship (but still most for chr1). Intermediate time points witnessed a progression between early and late patterns.

The changes over time suggested that Rec114 and Mer2 binding to chromosomes might be regulated by distinct mechanisms in early and late prophase, so we wished to obtain detailed information about association and dissociation from chromatin. To this end, we prepared a temporal summary at each Rec114 ChIP peak by fitting sigmoidal curves to the upward and downward slopes of the time courses of absolute ChIPseq signal, and defining the association and dissociation times as the points at which these fitted curves reached 50% of the maximum (see Methods).

These association and dissociation times were not uniform along chromosomes, instead showing large early and late domains (**Figure 1F**). Despite this internal fluctuation, Rec114 on average associated earlier and dissociated later from the shortest chromosome (chr1) compared to the longest (chr4) (**Figure 1F**) and this trend extended in a size-related manner across the entire complement (**Figure 1G**). Importantly, however, per-chromosome patterns differed between association and dissociation in a way that paralleled the ChIP density patterns noted above. Specifically, the three shortest chromosomes had especially precocious association times while the rest showed no clear size relationship (red points in **Figure 1G**). This mirrored relative ChIP densities at early times (**Figure 1E**, 2 h). In contrast, dissociation times showed a strong negative size correlation across all chromosomes, with the short trio fitting an overall linear relationship (blue points in **Figure 1G**). This mirrored the late ChIP density patterns (**Figure 1E**, 6 h).

We further estimated the average duration of Rec114 binding as the difference between association and dissociation times (green lines in **Figure 1F**). Mean Rec114 duration correlated negatively with chromosome size, and the short trio again stood out with especially long durations in all three datasets (**Figures 1H and S1D**). The degrees to which the shortest chromosomes showed early association and late dissociation, and hence long duration of Rec114 binding, were highly reproducible (**Figures 1I and S1E,F**). Mer2 showed highly similar patterns (**Figures S1C, S1D, and S1F**).

It is plausible that the binding duration of Rec114 and Mer2 (and probably other pro-DSB factors) dictates how long chromosomes are competent to make DSBs (Wojtasz et al., 2009) (Panizza et al., 2011; Carballo et al., 2013; Keeney et al., 2014). If so, then we would expect to see the Rec114 and Mer2 binding patterns reflected in DSB distributions. To test this, we mapped Spo11 oligonucleotides (oligos) at 4 and 6 h to compare the DSB landscapes at different times in prophase. Spo11 generates DSBs through a covalent protein-DNA complex that is endonucleolytically cleaved to release Spo11 bound to a short oligo (Keeney et al., 1997; Neale and Keeney, 2006). These oligos can be deep sequenced to generate DSB maps (Pan et al., 2011). Spo11-oligo densities at both time points were negatively correlated with chromosome size, as previously shown (Pan et al., 2011; Thacker et al., 2014) (**Figure S1G**). However, the short trio stood out as having substantially higher Spo11oligo density than predicted from a karyotype-wide linear relationship with chromosome size. Moreover, the 13 largest chromosomes showed a better negative correlation with size at 6 h than at 4 h. Thus, DSB distributions mirror the changes in Rec114 and Mer2 distributions.

We conclude that temporal regulation of DSB protein association and dissociation is likely to govern size-dependent DSB control on all chromosomes but also more specifically to ensure the necessary overrepresentation of DSBs and crossovers on very short chromosomes. We therefore set out to define the pathways that accomplish this temporal regulation.

### Distinct early and late mechanisms: homolog engagement dominates late patterns

DSB formation is inhibited when chromosomes successfully engage their homologs, and this negative feedback establishes an anticorrelation between DSB density and chromosome size in yeast (Kauppi et al., 2013; Keeney et al., 2014; Thacker et al., 2014). One hypothesis, supported by lifespans of resected DSBs (Mimitou et al., 2017), is that smaller chromosomes tend to terminate DSB formation later because they take longer on average to find their homologous partners (Keeney et al., 2014; Thacker et al., 2014). How homolog engagement inhibits DSB formation is not known, but studies in yeast and mice revealed that pro-DSB factors including Rec114 and Mer2 are preferentially bound to chromosome segments that have not synapsed with a partner (Kumar et al., 2010; Panizza et al., 2011; Kumar et al., 2015; Stanzione et al., 2016), and that formation of synaptonemal complex provokes active displacement of these proteins (Wojtasz et al., 2009; Carballo et al., 2013; Stanzione et al., 2016). A straightforward hypothesis is that feedback from homolog engagement explains the negative correlation between Rec114/Mer2 duration and chromosome size and thus also explains Rec114/Mer2 overrepresentation on small chromosomes.

However, in an earlier study documenting that Rec114 ChIP-chip signal is higher on the three short chromosomes at 4 h in meiosis, it was stated that the same Rec114 profile was found in a *spo11-Y135F* mutant, which cannot make DSBs (data not shown in (Panizza et al., 2011)). This seems to undermine the homolog engagement hypothesis: there is no homolog engagement without Spo11 activity, so there should be no difference between small and large chromosomes. Furthermore, we found striking DSB protein overrepresentation already at an early time point (2 h) (**Figures 1C, 1E, S1B, and S1C**), before synaptonemal complex is likely to have formed (Padmore et al., 1991; Henderson and Keeney, 2004) and before establishment of interhomolog recombination bias (Joshi et al., 2015).

We therefore considered an alternative, namely, that the chromosome size dependence of DSB protein binding is dominated by homolog engagement-mediated displacement late in prophase, but the early pattern is established by a distinct mechanism(s). This hypothesis predicts that disrupting homolog engagement should not affect the early pattern of Rec114 binding but should specifically disrupt the later pattern by causing Rec114 to be inappropriately retained on larger chromosomes.

To test this prediction, we performed Rec114 ChIP-seq in a mutant defective for homolog engagement (*zip3*) (Thacker et al., 2014). The *zip3* mutant had only modest differences in genome-average absolute Rec114 ChIP signal at 2 and 4 h (possibly reflecting small differences in culture timing), but had an ∼two-fold increase at 6 h, consistent with expectation if loss of homolog engagement causes Rec114 to persist longer than it should (**Figures 2A, and S2A**). More importantly, Rec114 was still overrepresented on the shortest chromosomes at 2 h and 4 h in *zip3*, with patterns similar to the same times in wild type (**Figures 2B, S2A and S2B**). At 6 h in contrast, the *zip3* mutant had failed to establish a strong negative correlation between ChIP density and chromosome size (**Figure 2B**), and this was because Rec114 was retained preferentially on larger chromosomes (**Figures 2C, S2A, and S2B**).

**Figure 2.**
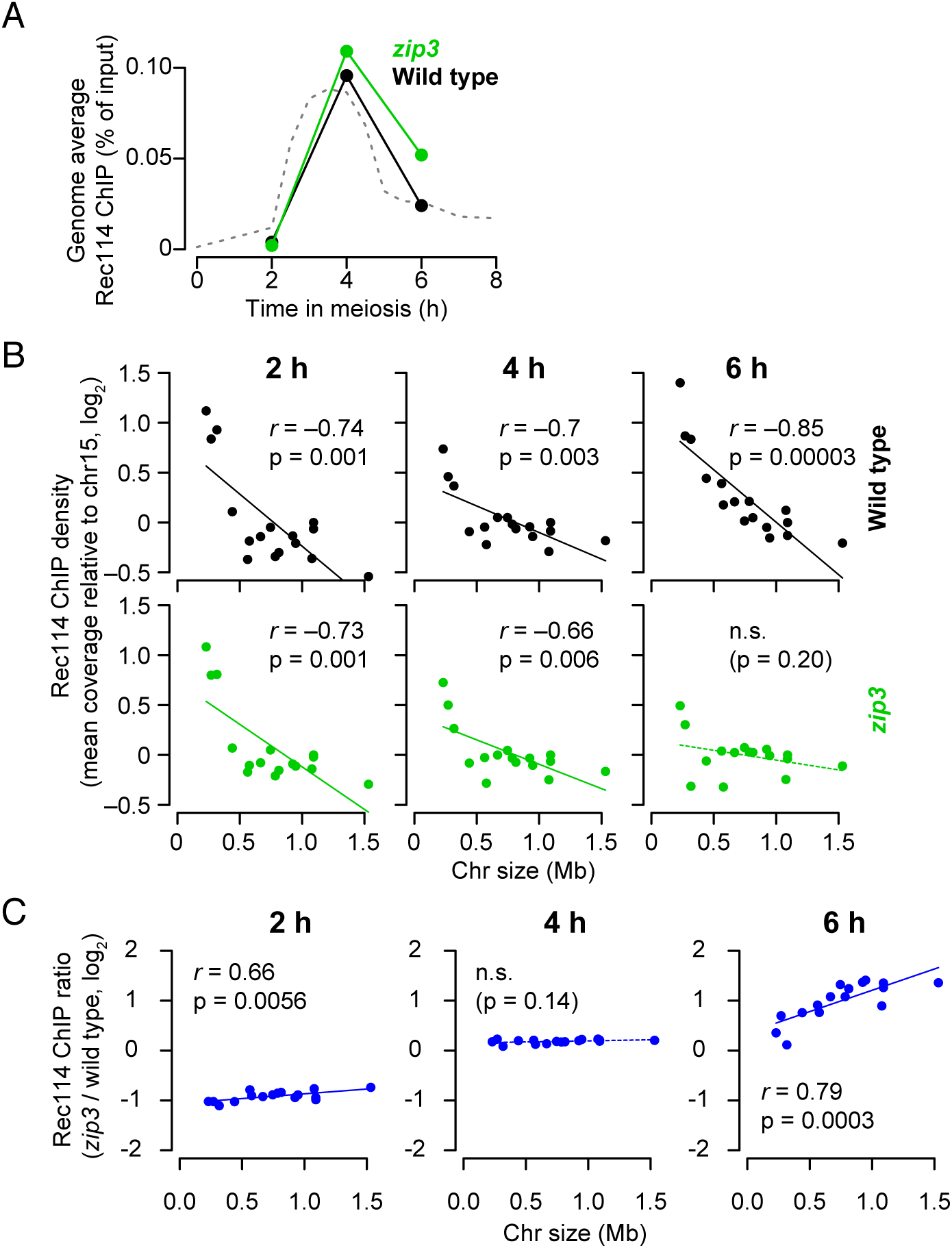
Zip3-dependent homolog engagement shapes Rec114 abundance late in prophase. (A) Genome-average Rec114 levels. Dashed line represents wild-type profile from Figure 1D. (B) Average per-chromosome Rec114 ChIP density (normalized to chr15). (C) The *zip3* mutation causes chromosome size-dependent changes in Rec114 distribution at 6 h but not earlier. See also Figure S2.

We conclude that feedback from homolog engagement establishes DSB control late in prophase by governing the dissociation of Rec114 and thus the duration of Rec114 binding to chromosomes. We further infer that the early patterns of Rec114 binding are shaped by a different mechanism(s), so we next set out to understand how the early patterns are shaped.

### Replication timing partially accounts for early Rec114 association on short chromosomes

Rec114 binding to chromatin is coordinated with meiotic replication (Murakami and Keeney, 2014). Thus, if short chromosomes replicate early, it might provide a straightforward explanation for their early Rec114 association. We tested this idea two ways, by examining the effect of replication delay caused by the origin deletions on chr3L, and by using the *tof1* mutant to compromise temporospatial coordination between replication and Rec114 recruitment (Murakami and Keeney, 2014). We used whole-genome sequence coverage from the ChIP input samples to measure relative replication times (“replication index”, i.e., log-fold difference in sequence coverage of a replicating sample compared to a premeiotic (non-replicating) sample).

In wild type, the three shortest chromosomes indeed exhibited relatively early replication, and chr3L naturally showed both very early replication and very early Rec114 association (**Figure 3Ai**). Origin deletion delayed both replication and Rec114 association on chr3L without affecting the right arm (chr3R), but, importantly, Rec114 association on chr3L was still on the early side when compared with longer chromosomes despite an extreme replication delay (**Figure 3Aii**). A similar pattern held when comparing Mer2 association time with replication index for the origin-deleted strain (**Figure S3A**). Moreover, in the *tof1* mutant the short trio still showed earlier Rec114 association than other chromosomes (**Figure 3Aiii**), and origin deletion delayed Rec114 association but to a lesser degree than it delayed replication (**Figure 3Aiv**).

**Figure 3.**
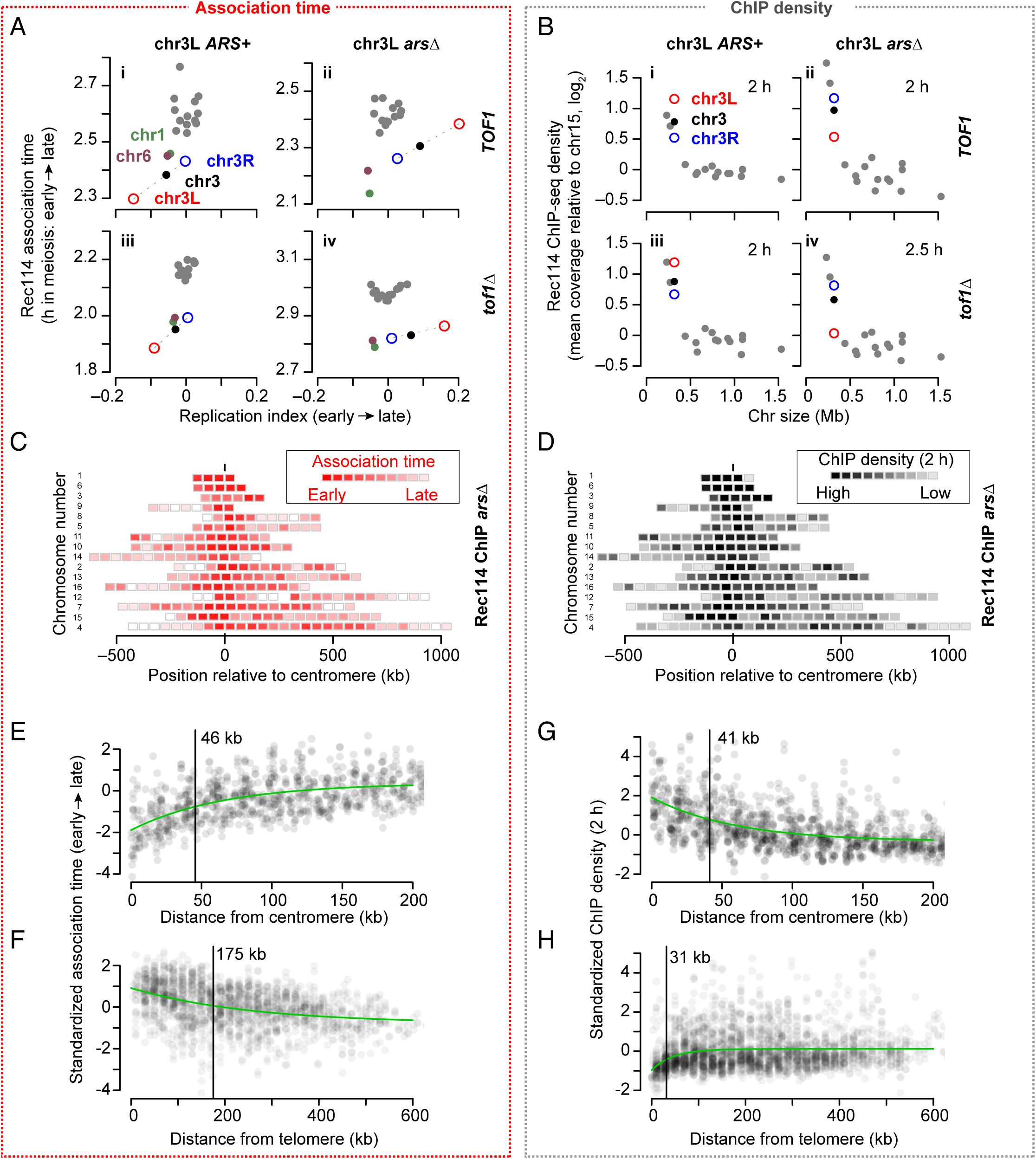
Replication timing and proximity to centromeres and telomeres influence time and amount of Rec114 association. (A) Comparison of per-chromosome Rec114 association time with replication timing. Replication index is defined as –log2 of the ratio of sequence coverages for samples in S phase vs. G1 phase; lower values indicate earlier replication (Murakami and Keeney, 2014). The means for the left arm (chr3L, open red circle) and right arm (chr3R, open blue circle) of chr3 are plotted separately, connected to the means for the entire chromosome (solid black) by dashed lines. (B) Average per-chromosome Rec114 ChIP density at 2 h (normalized to chr15). Left and right arms of chr3 are displayed separately along with the whole-chromosome mean. (C,D) Intra-chromosomal distributions of Rec114 association times (C) and ChIP density at 2 h (D) in the *arsΔ* strain. Each block represents a 50-kb bin color-coded according to the average of the Rec114 association times for peak positions within the bin, or average ChIP density for the bin. Chromosomes are ranked by size, with centromere at position zero. Other datasets are shown in Figures S3C and S3D). (E–H) Centromere and telomere effects on association time (E, F) and ChIP density (G, H) of DSB proteins. The two Rec114 and one Mer2 ChIP time courses were combined as follows. To capture the intra-chromosomal features separate from inter-chromosomal difference, we averaged each dataset in 20-kb bins, standardized the values for each chromosome to a mean of 0 and variance of 1, and standardized again within a dataset before pooling all datasets together. The first standardization minimizes differences between chromosomes and the second standardization minimizes differences between datasets. Each point is the value from one dataset for a 20-kb bin, plotted as a function of distance from centromere (E, G) or telomere (F, H). Green lines are fitted exponential models. Vertical bars indicate the distance where the effect decays to half of the original value. See also Figure S3.

The early density of Rec114 ChIP signal exhibited a complementary trend. Rec114 was naturally overrepresented on chr3L (**Figure 3Bi**). Origin deletion caused a substantial decrease on chr3L but still left Rec114 at higher levels than on the larger chromosomes (**Figure 3Bii**) and Mer2 showed a similar pattern (**Figure S3B**). In the *tof1* mutant, the short trio still showed high Rec114 signal (**Figure 3Biii**), and origin deletion reduced this signal on chr3L but left it in the higher part of the longer-chromosome range (**Figure 3Biv**).

These findings indicate that coordination with replication timing promotes early association and preferential recruitment of Rec114 and Mer2 on short chromosomes, but is not sufficient to explain all of the differences from longer chromosomes. We thus infer that additional control layers exist.

### DSB proteins accumulate earlier near centromeres and later near telomeres

To further understand the control of DSB protein behavior, we examined the domain structure noted above, color-coding 50-kb bins according to their association times or ChIP densities for Rec114 and Mer2 (**Figures 3C, 3D, S3C and S3D**). These heatmaps suggested a global tendency of early association and higher ChIP densities around centromeres (originally suggested by (Kugou et al., 2009)), and the converse toward chromosome ends. To systematically quantify these spatial patterns, we combined the datasets, plotted all 32 chromosome arms together as a function of distance from centromere or telomere, and fitted exponential trend lines (**Figures 3E–H**).

As we surmised from the heatmaps, DSB proteins tended to show early association near centromeres (**Figure 3E**). This effect decreased with distance, reaching half its initial strength 46 kb away. Conversely, telomere-proximal regions showed late association, and this effect decayed to half strength at 175 kb (**Figure 3F**). This telomere effect may be related to a previously described DSB formation delay (Borde et al., 2000). Centromere and telomere effects were both retained in *tof1* mu-tants, but appeared weaker (**Figure S3E**), possibly indicating that both effects depend partially on coupling of DSB protein association to constitutively early or late replication in centromere- and telomere-proximal regions (Blitzblau et al., 2012), respectively.

Complementary patterns were again seen when we analyzed early ChIP density (at 2 h). Centromere-proximal regions had higher ChIP density and this effect decayed by half at 41 kb (**Figure 3G**), while telomere-proximal regions had lower ChIP density, with effect half-maximal at 31 kb (**Figure 3H**). The similar shapes of trend lines for peri-centromeric association time and ChIP density might indicate that both features of DSB protein binding reflect the same underlying mechanism. The Mer2 dataset had more samples for early time points, affording a detailed look at early prophase. Pericentromeric enrichment of Mer2 was detectable at 0.5 h in meiosis, reached a maximum at 1.5 h, then gradually disappeared as Mer2 accumulation elsewhere balanced out binding near centromeres (**Figure S3F**). In contrast, there was little or no sign of telomere-proximal depletion of Mer2 through 1 h, then it became progressively stronger thereafter (**Figure S3G**). The centromere effect was normal in the *zip3* mutant (**Figure S3H**), as expected if homolog engagement influences late but not early patterns of DSB protein binding.

### A model integrating replication, centromere, and telomere effects explains all but the shortest chromosomes well

To test if a combination of replication timing and the centromere and telomere effects might be sufficient to account for Rec114 association timing, we built an integrative model using multiple linear regression. Replication timing and the centromere effect were correlated with Rec114 association timing (*r* = 0.49 and 0.66, respectively) while the telomere effect was less so (*r* = 0.23) (**Figure S4A**). Replication timing and the centromere effect correlated with each other (*r* = 0.47) because centromere-proximal regions tend to replicate early (Blitzblau et al., 2012), whereas the telomere effect was not correlated with the other variables.

Multiple regression models with replication timing and the centromere and telomere effects as the explanatory variables accounted for 37% and 51% of the variance in Rec114 association timing (*ARS+* and *arsΔ*, respectively; **Figure 4A**) and 40% for Mer2 association (**Figure S4B**). Thus, this simple three-factor model is reasonably effective at fitting Rec114 association genome-wide (**Figure 4B**). Strikingly however, the model fit the shortest chromosomes poorly, with observed association times consistently earlier than predicted by the model (**Figures 4A, 4C and S4B**). ChIP density at 2 h gave complementary findings: the models fit genome-wide data well but consistently underperformed on the smallest chromosomes by predicting less relative enrichment of Rec114 or Mer2 than was observed (**Figure 4D–F and S4B**).

**Figure 4.**
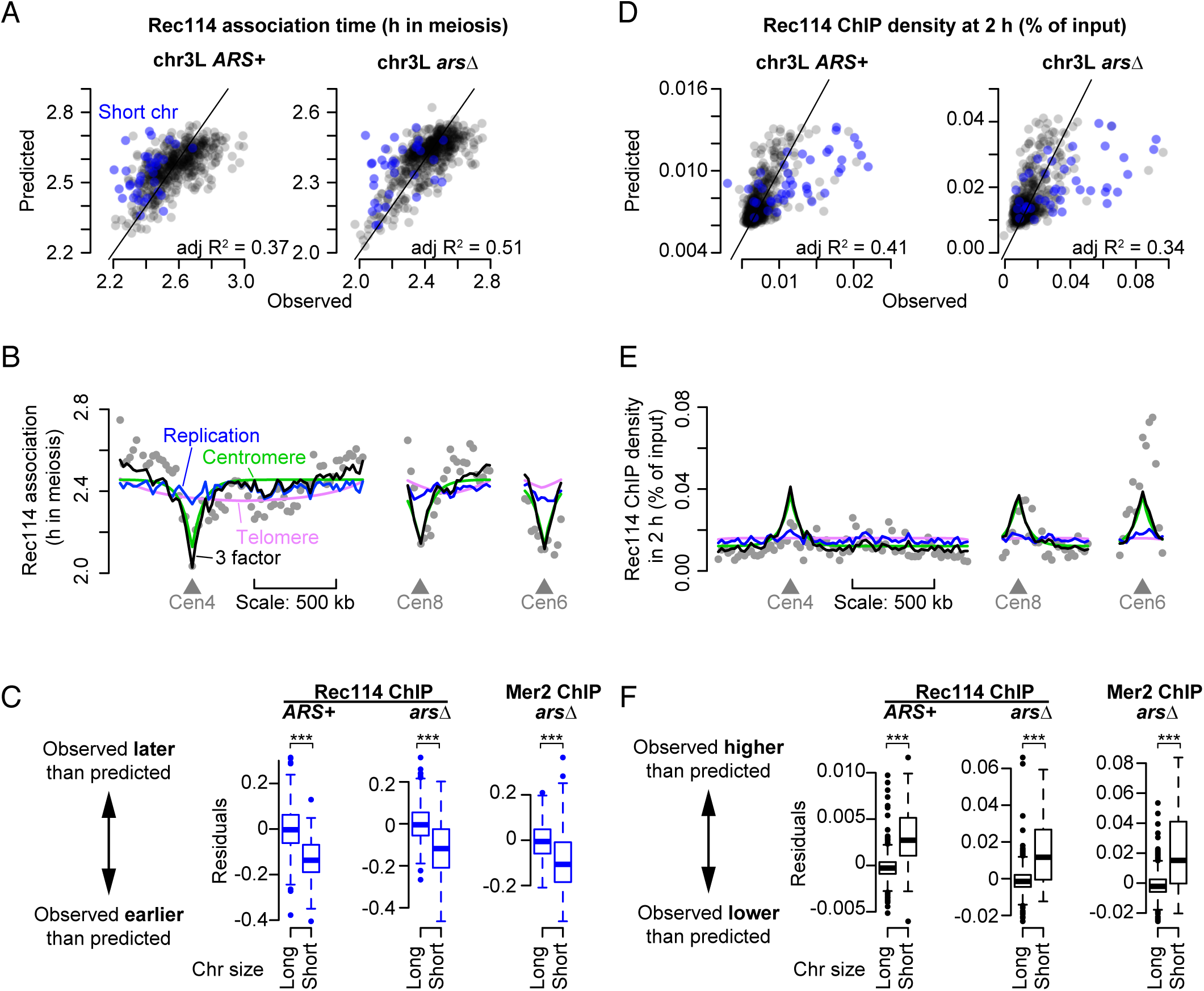
DSB protein binding is boosted on the smallest chromosomes early in meiosis. (A,D) A model incorporating replication timing and the centromere and telomere effects. Replication index and centromere and telomere effects were binned in 20-kb windows and used as explanatory variables to model Rec114 association time (A) or ChIP density at 2 h (D) by multiple linear regression. Each point compares the observed and model-predicted value for Rec114 association time within a bin. Points for bins on short chromosomes are blue. (B, E) Examples of within-chromosome patterns predicted by the multiple regression model and each of its component factors. Gray dots are averages Rec114 association times (B) or Rec114 ChIP density at 2 h in 20-kb bins (E, *arsΔ* dataset). Blue, green, and magenta lines are the component replication, centromere, and telomere effects, respectively, and the black line is the prediction from the three-factor regression model. (C,F) Multiple regression models systematically underperform on the short chromosomes. The three-factor multiple regression models were applied to each dataset in turn and the residuals were grouped for the three shortest chromosomes vs. the rest except chr12. *** indicates p < 0.005, one-sided Wilcoxon test. See also Figure S4.

These findings suggest that, for most chromosomes, DSB protein association early in meiotic prophase is strongly shaped by the composite influences of replication timing and proximity to centromere or telomere. The shortest trio, in contrast, accumulates DSB proteins earlier and at higher levels than these composite influences predict. We thus hypothesized that the little chromosomes have an additional feature(s) that boosts their ability to compete for DSB protein binding early. This idea makes two simple predictions if this feature is intrinsic to the DNA sequence: segments from the short chromosomes should retain the boost even when fused to a longer chromosome, but making an artificially small chromosome by bisecting a larger one should not be sufficient to establish a similar boost. As detailed next, both predictions were met.

### Intrinsic features of short chromosomes promote preferential Rec114 binding

To test if chr1 (the smallest at 230 kb) has an intrinsic feature(s) that boosts Rec114 binding, we placed it in an artificially lengthened context by targeting a reciprocal translocation with chr4 (at 1.5 Mbp, the longest chromosome if we exclude the rDNA array on chr12) (**Figures 5A, S5A and S5B**). We then asked whether the chr1-syntenic portions of the derivative chromosomes (der(1) at 532 kb and der(4) at 1.2 Mb) still behaved like one of the shortest chromosomes as our hypothesis predicts (i.e., high-level Rec114 binding at early times) or if instead they now behaved like a longer chromosome as previous studies of crossing over on similar translocation chromosomes might predict (Kaback et al., 1992).

**Figure 5.**
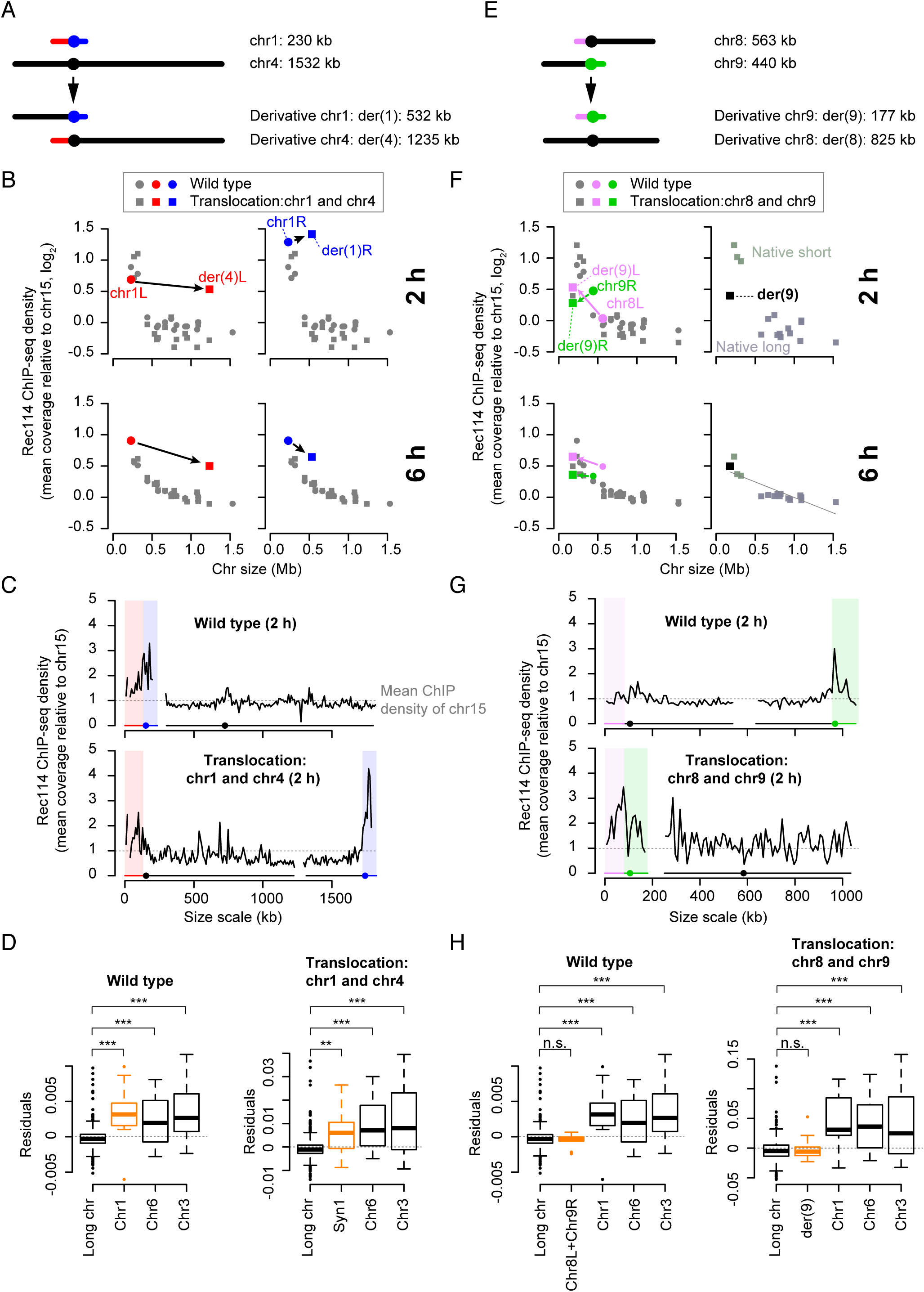
The Rec114 boost is intrinsic to short chromosomes. (A) Targeted translocation between chr1 and chr4 (to scale). (B) Chr1-derived sequences retain high-level Rec114 binding when in a large-chromosome context. Average per-chromosome Rec114 ChIP densities normalized to chr15 are shown at 2 and 6 h. (C) Rec114 profiles for wild-type and translocated chromosomes. ChIP-seq data were normalized relative to chr15 and smoothed with a 10-kb sliding window. (D) A three-factor multiple regression underperforms on chr1-derived sequences in both wild-type and translocation contexts. Boxplots show residuals from multiple regression performed as in Figure 4D using the 2 h data. “Syn1” indicates sequence syntenic to chr1. (E) Targeted translocation between chr8 and chr9 (to scale). (F–H) An artificially short chromosome fails to acquire a boost in Rec114 binding. Per-chromosome average Rec114 ChIP densities (F), Rec114 profiles (G), and multiple regression residuals (H) are shown as for panels B, C, and D, respectively. In panels D and H, ***: p < 0.005; **: p < 0.01; *: p ≤ 0.05; and n.s.: not significant (p > 0.05), one-sided Wilcoxon test. See also Figure S5.

In wild type at 2 h, both arms of chr1 exhibited the characteristic Rec114 overrepresentation, and this preferential Rec114 binding was fully retained for the chr1-syntenic parts of der(1) and der(4) (**Figures 5B, upper graphs**). As a result, there was a sharp transition at the boundaries between chr4- and chr1-derived sequences in the ChIP profiles (**Figure 5C**). Moreover, a three-factor multiple regression model again underperformed in predicting Rec114 levels specifically on the chr1-derived sequences (plus chr3 and chr6), whether in the native context or the translocation with chr4 (**Figure 5D**). These results strongly support the hypothesis that chr1 has an intrinsic feature(s) that promotes preferential Rec114 association at an early stage of meiotic prophase, and further show that its effects are limited in cis and independent of chromosome size per se.

Interestingly, at later time points (4 h and especially 6 h), segments from chr1 still showed Rec114 overrepresentation in the translocation context, but to a lesser degree than native chr1 (**Figure 5B, lower graphs, and Figures S5C and S5D**). This reduced Rec114 abundance matches expectation from our conclusion that feedback from homolog engagement is a dominant factor late in prophase. Since homolog engagement is tied to chromosome size per se, we expect the chr1-derived segments to behave in line with their long chromosomal context. More specifically, we anticipate that they would tend to engage their homologs earlier and more efficiently on average than in wild type, and thus to dissociate Rec114 earlier. The remaining degree of overrepresentation relative to naturally long chromosomes may be a residuum of the exceptionally high enrichment seen earlier. A recently described tendency of telomere-adjacent regions to retain Hop1 at late times may contribute to Rec114 retention as well (Subramanian et al., 2018).

To carry out the converse experiment, we created an artificial short chromosome by engineering a translocation between two medium-size chromosomes (**Figures 5E**, **S5A and S5B**). The smaller of the derivative chromosomes (der(9), 177 kb) contained the left arm of chr8 (now der(9)L) and the right arm of chr9 (now der(9)R). If our supposition is correct that the shortest chromosomes benefit from an intrinsic boost that does not occur on these segments of chr8 and chr9, then the artificial short chromosome should still be fit well by the regression model.

At 2 h, relative Rec114 ChIP density was ∼40% higher on der(9)L but slightly lower on der(9)R as compared with the same segments in their normal contexts (**Figure 5F, upper left, and Figure 5G**). The net effect was a modest elevation in chromosome-average Rec114 signal for der(9) relative to the native long chromosomes, but substantially below the native short trio (**Figure 5F, upper right**). Moreover, a three-factor regression model still predicted Rec114 levels very well on der(9) (note the small residuals in **Figure 5H**). These findings support the conclusion that these segments of chr8 and chr9 do not share the apparent Rec114binding boost seen on the shortest trio.

Relative Rec114 ChIP density was also elevated on der(9) at 4 h and 6 h, but importantly, the degree of elevation was now closely in line with that on the shortest trio (**Figures 5F, lower panels, and S5E and S5F**). Thus, for late Rec114 binding patterns, the artificial short chromosome behaved like a natural short chromosome. This result is readily understood in light of the homolog engagement model, if one assumes that der(9) establishes homolog engagement more slowly and/or less efficiently than either chr8 or chr9.

### Evolutionary selection maintains preferential Rec114 recruitment

Most species in the *Saccharomyces sensu stricto* complex have the same three very short chromosomes (Fischer et al., 2000). The remarkable evolutionary stability of this potentially risky karyotype suggests that mechanisms mitigating the risk of meiotic nondisjunction are shared across the ∼20 Myr since the last common ancestor of this clade, implying in turn the existence of selective pressure to maintain hyperrecombinogenic properties of the smallest chromosomes.

One member of this clade, *Saccharomyces mikatae*, provides a natural experiment to test for hallmarks of such selective pressure. This species contains natural translocations between ancestral chr6 and chr7 (Fischer et al., 2000; Kellis et al., 2003). In other *Saccharomyces* species, chr6 is the second shortest chromosome, but in *S. mikatae* the regions syntenic to *S. cerevisiae* chr6 (hereafter, syn6) are on longer chromosomes (**Figure 6A**). We previously showed that syn6 regions have a DSB density matching their chromosomal context (i.e., low in *S. mikatae* relative to other species) from which we inferred that whole-chromosome DSB density is tied to chromosome size per se and is largely extrinsic to the DNA sequence (Lam and Keeney, 2015). We were prompted to revisit this conclusion in light of our new findings of multilayered control of DSB potential and the contribution of intrinsic features of the smallest chromosomes.

**Figure 6.**
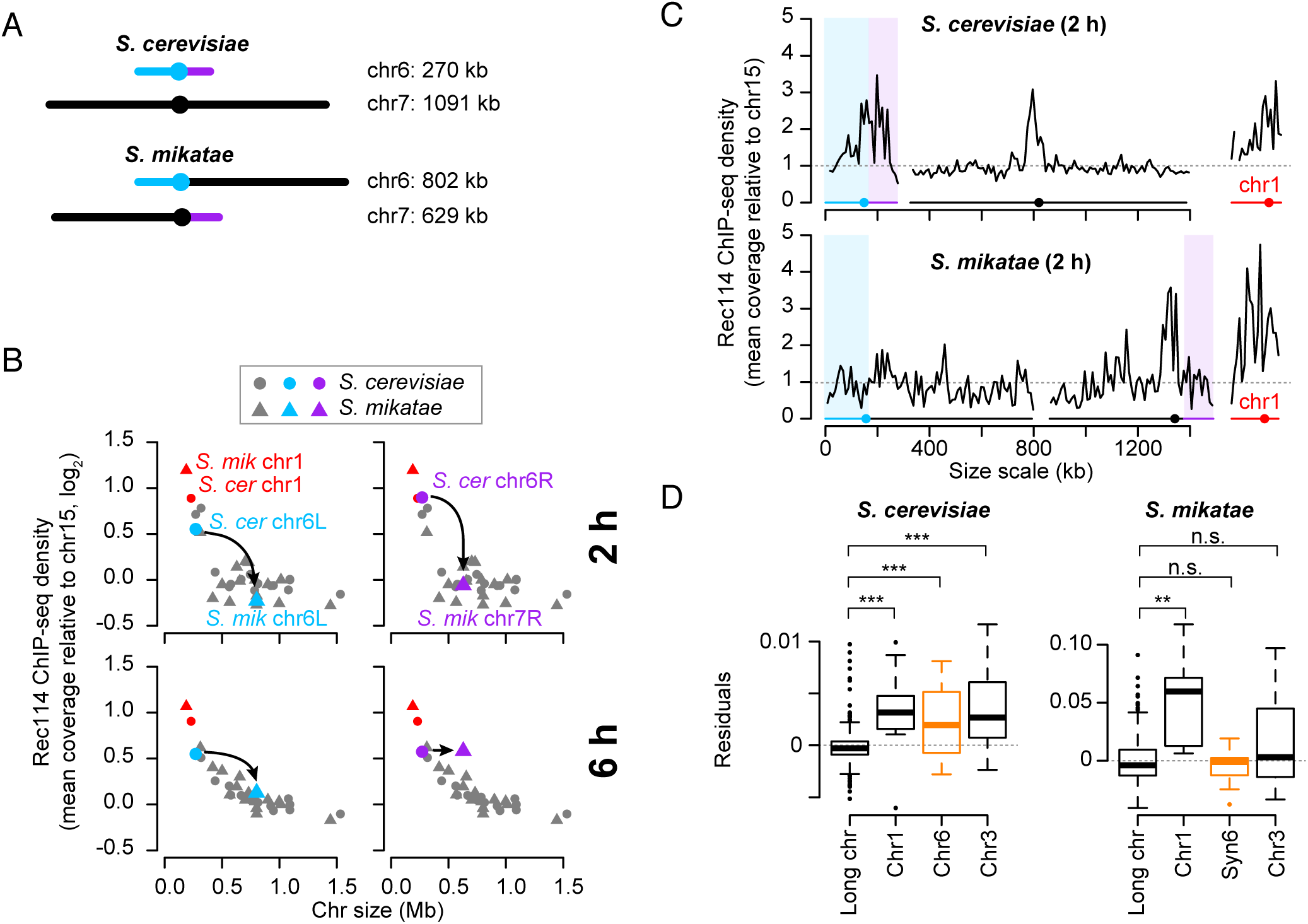
Selective pressure has maintained the short chromosome boost on chr6. (A) Syntenic regions that reside on different-sized chromosomes in *S. cerevisiae* and *S. mikatae* (to scale). (B–D) Regions syntenic to ancestral chr6 do not show a Rec114 binding boost in *S. mikatae*. Per-chromosome average Rec114 ChIP densities (B), Rec114 profiles (C), and multiple regression residuals (D) are shown as for Figure 5, panels B, C, and D, respectively. Note that the model also underperformed for chr3 in *S. mikatae*, but the distribution of residuals was not statistically significant (p = 0.077). See also Figure S6.

We reasoned that, if there is selective pressure to maintain a DSB protein-binding boost on the smallest chromosomes, then this selective pressure would have been absent for syn6 regions during the posttranslocation generations of the *S. mikatae* lineage. Without selection to maintain it, the boost might decay and thus be attenuated or absent in extant members of this species. Conversely, selective pressure should have preserved the boost on chr1 because it is still small. To test these predictions, we performed Rec114 ChIP-seq on meiotic *S. mikatae* cultures.

As we predicted, relative Rec114 binding at 2 h was much lower on syn6 in *S. mikatae* (chr6L and chr7R in that species) than for chr6 in *S. cerevisiae*, such that syn6 segments were indistinguishable from any other segment on a medium or large chromosome (**Figure 6B, upper graphs, and 6C**). In contrast, chr1 in *S. mikatae* showed strong overrepresentation of Rec114 (**Figures 6B and 6C**). Furthermore, a three-factor regression model fit the data for syn6 well in *S. mikatae*, but showed the characteristic underperformance for chr1 (**Figure 6D**). At later time points (4 and 6 h), syn6 segments had relative Rec114 densities more in line with their chromosome sizes (**Figure 6B, lower graphs, S6A and S6B**). Collectively, these findings support the hypothesis that the smallest chromosomes in *Saccharomyces* species are under selective pressure to maintain exceptionally high-level DSB protein binding.

### Chromosome axis proteins are required for the short chromosome boost

How do the smallest chromosomes preferentially recruit DSB proteins? To begin to address this question, we examined contributions of higher order chromosome structures. Meiotic chromosomes form proteinaceous axial structures from which chromatin loops emerge (Kleckner, 2006). Several proteins involved in forming these axes (Hop1, Red1, and Rec8) are also important for normal chromatin association of Rec114 and Mer2 (Panizza et al., 2011). Hop1 and Red1 themselves are overrepresented on short chromosomes in a mutually dependent manner, so it was proposed that axis protein enrichment contributes to the high density of DSBs on the smallest chromosomes (Panizza et al., 2011; Sun et al., 2015). Moreover, preferential enrichment of axis proteins may support enrichment of DSB proteins. To test these hypotheses, we assessed Rec114 ChIP density and DSB formation in axis protein mutants.

Both *hop1* and *red1* single mutants eliminated Rec114 overrepresentation on the smallest chromosomes early in prophase (2 h) and also essentially ablated the strong negative dependence of Rec114 binding on chromosome size at all times assayed (**Figure S7A**). However, both mutations greatly decreased Rec114 ChIP levels genome-wide (Panizza et al., 2011) (**Figure 7A**), raising the concern that low signal:noise ratio in these mutants might obscure true chromosome-specific patterns. To rule out this possibility, we took advantage of our observation that Rec114 chromatin binding is substantially higher in a *hop1 red1* double mutant than in the single mutants (**Figure 7A**), possibly because of negative effects of residual binding of Hop1 or Red1 in the absence of the other (Sun et al., 2015). Importantly, even with this more robust ChIP signal, the shortest chromosomes still failed to show Rec114 overrepresentation at 2 h, and only a very weak chromosome size dependence emerged at later times (**Figure 7B**).

**Figure 7.**
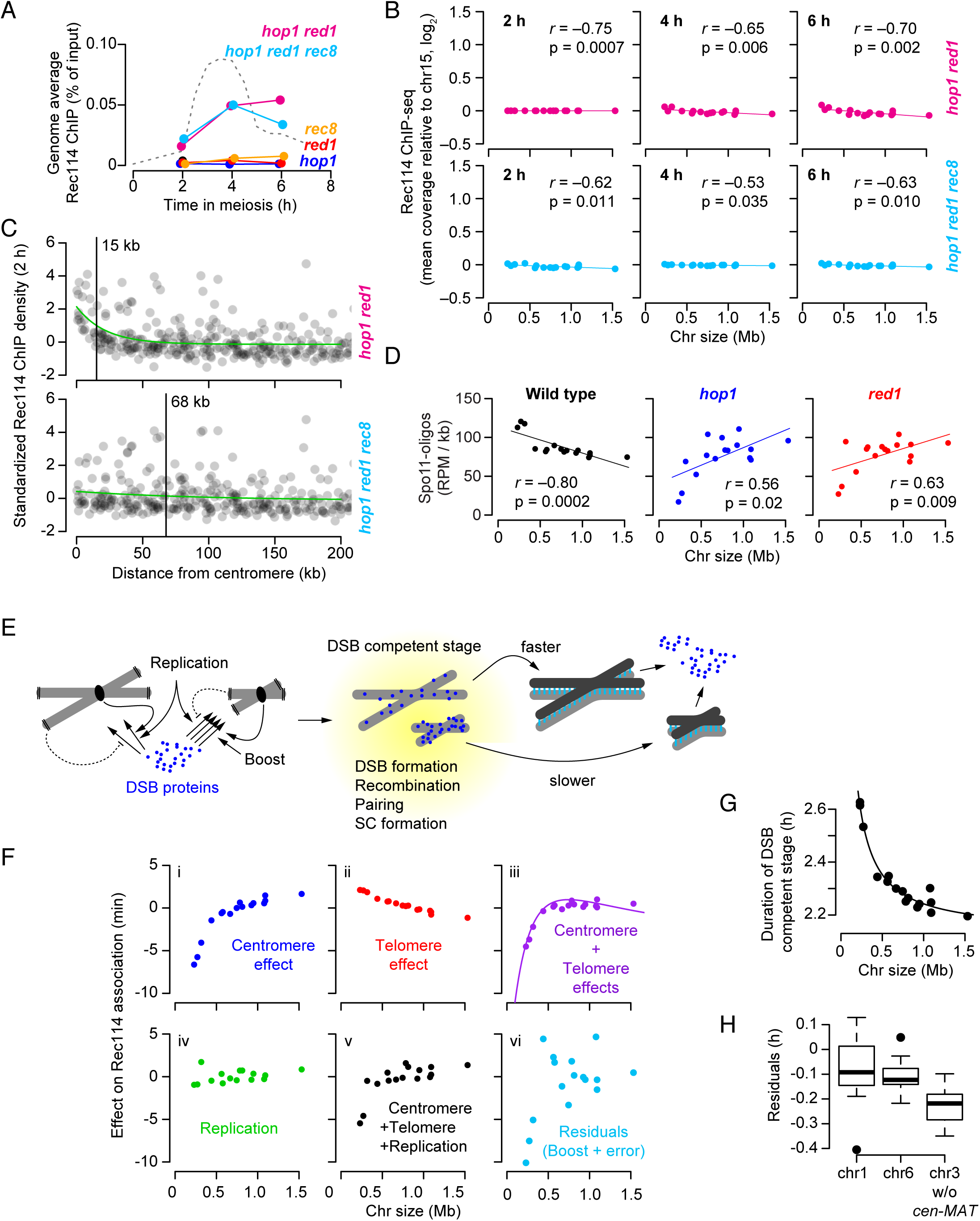
Axis proteins are required for the short chromosome boost. (A) Genome-average Rec114 levels. Dashed line represents wild-type profile from Figure 1D. (B) Chromosome size dependence of Rec114 binding to chromosomes is lost in the absence of Hop1 and Red1. Note that, although correlations with chromosome size are statistically significant, their slopes are negligible compared to wild type (Figure 1E). (c) The centromere effect is lost in the absence of Rec8 but still present (albeit spreading less far) in the absence of Hop1 and Red1. Rec114 ChIP data at 2 h were binned and standardized as described for Figure 3E. (D) Preferential DSB formation on small chromosomes is lost in hop1 and red1 mutants. (E) Cartoon depicting the multiple pathways that govern DSB protein association and dissociation from long vs. short chromosomes. (F) Size dependence of the contributions of the individual early pathways and their integration. Parameters are estimated from the three-factor model fitted to Rec114 association time in the arsΔ strain. Panels i and ii estimate the net effect of the centromere and telomere effects on each chromosome’s Rec114 association time and panel iii shows their combined effect. Panel iv estimates the per-chromosome effect of replication timing (note that chr3 is an outlier because of the arsΔ on the left arm), and panel v merges effects of all three pathways. Panel vi depicts what is not explained by these three pathways, inferred as the effect of the short chromosome boost. (G) The per-chromosome duration of DSB protein binding follows an inverse proportion relationship with chromosome size. Duration data from Figures 1H and S1D were combined; the cold region between *CEN3* and *MAT* was censored. (H) Residuals for each member of the shortest chromosome trio (not including *CEN3–MAT* interval) from the three-factor multiple regression model applied to Rec114 association time (*ARS+* dataset, Figure 1C). See also Figure S7.

Absence of the meiotic cohesin subunit Rec8 does not eliminate enrichment of Red1 on the shortest chromosomes (Sun et al., 2015). Similarly, we found that Rec114 overrepresentation still occurred on the short trio in a *rec8* mutant (**Figure S7A**), even though total Rec114 levels were greatly reduced (**Figure 7A**). A *hop1 red1 rec8* triple mutant behaved like a *hop1 red1* double mutant (improved total Rec114 recruitment but no short-chromosome boost) (**Figures 7A and 7B**). Given that Rec8 is particularly enriched around centromeres (Blat et al., 2002; Kugou et al., 2009), we also examined its potential role in the preferential binding of Rec114 near centromeres. *rec8* mutants no longer showed higher Rec114 binding around centromeres (**Figure S7B**). The centromere effect was retained in *hop1* and *red1* single and double mutants, albeit extending less far (**Figures 7C and S7B**). The *rec8* mutation was epistatic to *hop1 red1* for loss of the centromere effect (**Figure 7C and S7B**).

Spo11-oligo maps demonstrated the functional significance of loss of Hop1 and Red1 for the short-chromosome boost: the axis mutants lacked a markedly higher density of DSBs on the shortest chromosomes at 4 h (**Figure 7D**). Instead, there was now an inverted relationship between DSB density and chromosome size in the *hop1* and *red1* mutants, indicating a complete loss of size-dependent DSB control.

Taken together, these findings indicate that Hop1 and Red1 but not Rec8 are critical for the short chromosome boost of both Rec114 binding and DSB formation. Conversely, Rec8 promotes Rec114 recruitment near centromeres while Hop1 and Red1 are largely dispensable. The findings further indicate that components of the meiotic axis are also needed for normal establishment of later chromosome size-related patterns of DSB protein enrichment, implicating the chromosome axis as a platform supporting most if not all features of DSB regulation.

## Discussion

We sought to understand how cells control DSB formation on a per-chromosome basis, focusing on when and where proteins essential for DSBs bind to chromatin (**Figure 7E**). Three distinct mechanisms govern spatial and temporal patterns of Rec114 and Mer2 chromatin association early in prophase on all chromosomes: replication timing, distance to the centromere, and distance to telomeres. We also identified a fourth early mechanism(s) that boosts DSB protein binding specifically on the three smallest chromosomes. We further found that yet another pathway (homolog engagement) controls duration of a DSB-permissive state by regulating DSB protein dissociation from chromosomes. These five pathways (and perhaps more) collaborate to shape the per-chromosome and within-chromosome DSB landscape, and to ensure that every chromosome has the opportunity to pair and recombine.

### Integrating multiple pathways with distinct chromosome size dependencies

To better understand the early pathways, we estimated their absolute effects on Rec114 association time (**Figure 7F**). Because centromere proximity speeds up Rec114 association while telomere proximity delays it, and because these pathways’ strengths decay exponentially on different scales, their chromosome size dependencies differ substantially. The centromere effect is highly non-linear with size, disproportionately influencing the three smallest chromosomes and giving a weaker linear size correlation on the remaining thirteen (**Figure 7Fi**). In contrast, the telomere effect is linearly anticorrelated with size across all chromosomes: the smallest chromosomes are delayed the most but not disproportionately so (**Figure 7Fii**). Combining these opposing effects leaves the shortest trio with relatively accelerated Rec114 association while the rest of the chromosomes show little difference among themselves (**Figure 7Fiii**). Replication timing correlates modestly with chromosome size (**Figure 7Fiv**), so adding this effect reinforces the early advantage of the smallest chromosomes and leaves only a weak size relationship on the larger chromosomes (**Figure 7Fv**).

We initially inferred existence of a small chromosome boost because Rec114 association was not fully explained by the combination of replication timing and the centromere and telomere effects (**Figure 7Fvi**). Overall, the small-chromosome boost and (to a lesser extent) the centromere effect contribute the most to privileging Rec114 binding on the smallest trio. Replication timing contributes more modestly, while telomeres have a small counter-balancing effect. It is unclear if the boost is a single mechanism or multiple, or if all three short chromosomes share the same mechanism. If multiple mechanisms are involved, they share a requirement for Hop1 and Red1. The chr1 translocations maintained the boost over one round of meiosis and multiple rounds of mitotic division during strain construction, so we infer that the boost relies on characteristics embedded in the chromosomal DNA sequence, not solely on epigenetic factors.

The homolog engagement pathway defines when pro-DSB factors are removed from chromosomes, and thus dictates the per-chromosome average Rec114 duration. It was previously proposed that the size dependence of homolog engagement is because the speed of homologous pairing and synapsis is governed by the number (not density) of DSBs (Keeney et al., 2014; Thacker et al., 2014; Mimitou et al., 2017). Hochwagen and colleagues suggested a nonexclusive alternative based on their finding that ∼100-kb long regions near telomeres of most chromosomes are resistant to synapsis-dependent downregulation of DSB formation (Subramanian et al., 2018). They proposed that the chromosome size dependence of homolog engagement arises because these telomere-associated regions are a larger fraction of smaller chromosomes (Subramanian et al., 2018). Both models predict an inverse proportional relationship to chromosome size, and indeed a simple inverse proportion model effectively described the chromosome size dependence of DSB protein duration (**Figure 7G**). The net effect of homolog engagement is thus to reinforce the nonlinear relationship between DSB protein binding and chromosome size, such that the smallest trio is highly privileged and the remaining thirteen show a more linear relationship with size (**Figure 7G**).

An important difference between the four early pathways and homolog engagement is that the early pathways proactively establish when chromosomal segments become competent to make DSBs. As such, these pathways principally modulate the population-average DSB probability. In contrast, homolog engagement is reactive because it acts on a per-cell and per-chromosome basis in response to a favorable outcome (presumably successful pairing, SC formation, and/or crossover designation). Thus, homolog engagement does not simply bias the population-average DSB probability in favor of smaller chromosomes (which it does do), it also actively ensures a low failure rate by allowing DSB formation to proceed until success is achieved. Other reactive regulatory circuits involving Tel1/ATM and Mec1-dependent control of pachytene exit operate in parallel to further ensure success (Lange et al., 2011; Zhang et al., 2011; Carballo et al., 2013; Gray et al., 2013; Cooper et al., 2014; Keeney et al., 2014).

### Establishing and realizing DSB potential

Cells establish “DSB potential” through control of the timing of DSB-promoting factors on chromosomes, but the amount of DSB proteins is probably also important. It is also likely that control of amount and timing are related but separable. For example, we envision that higher affinity binding sites outcompete weaker ones at early times when DSB proteins are first induced and at limiting concentrations, so high affinity sites will also tend to be earlier ones. Conversely, segments that are early for other reasons (e.g., proximity to early-firing replication origins) may not necessarily accumulate the highest levels of Rec114 etc., but would have a head start over other regions.

Once established, DSB potential must be realized in the form of DSBs and, eventually, recombination. Just as DSB potential is the net result of multiple integrated pathways, so too are DSB and recombination frequencies the net result of many influences. A good example of this complexity is provided by pericentromeric regions, where multiple, often mutually antagonistic, pathways work on different size scales to control DSB formation and recombination. Crossovers near centromeres can provoke missegregation (Rockmill et al., 2006), so local DSB protein enrichment could be deleterious. However, centromere-bound kinetochore components suppress DSB formation at short range (∼6-kb scale) and Rec8 inhibits interhomolog recombination for DSBs within ∼20 to 50 kb of centromeres (Vincenten et al., 2015). Interestingly, suppression of interhomolog recombination and centromeric enrichment of Rec114 work on similar size scales and both require Rec8, suggesting they may be mechanistically related.

### Karyotype constrains the DSB landscape in yeast and beyond

Our findings demonstrate how complex mechanisms collaborate to mitigate the missegregation risk of the smallest chromosomes. An alternative could be to avoid having short chromosomes altogether, but the apparently risky karyotype of *S. cerevisiae* is an ancient and evolutionarily successful one. This stability may imply that having these short chromosomes carries a fitness benefit that remains to be elucidated.

Chr3 has additional characteristics that shape its DSB landscape. Recombination is lower than genome average in the 86 kb from the centromere (*CEN3*) to the mating type locus (*MAT*), mostly because this is a cold zone for DSB formation (Baudat and Nicolas, 1997). It has been argued that linkage of *MAT* to *CEN3* favors maintenance of heterozygosity for centromere-linked genes after mating of spores from the same tetrad (automixis), and/or promotes formation of spores of the opposite mating type when sporulation produces a dyad instead of a tetrad under nutrient limitation (Taxis et al., 2005; Knop, 2006; Lacefield and Ingolia, 2006). Thus, it appears that pressure to maintain *CEN3–MAT* linkage has selected for suppression of DSB formation across this region throughout the *Saccharomyces* clade (Lam and Keeney, 2015). A further consequence is that DSB frequencies are elevated even more in the flanking regions to compensate for low DSB frequency between *CEN3* and *MAT*. This compensation involves high-level recruitment of axis and DSB proteins (Blat et al., 2002; Panizza et al., 2011) (this study). Indeed, the three-factor regression model underperformed to an even greater extent on the parts of chr3 excluding *CEN3– MAT* than it did for chr1 and chr6 (**Figure 7H**), indicative of an even stronger boost of Rec114 binding.

Boosting DSB protein binding is not unique to yeast. This strategy is also used in mammals to accomplish a similar end, i.e., ensuring high-level DSB formation in small chromosome segments that would otherwise risk missegregation. In most placental mammals, accurate segregation of the sex chromosomes in male meiosis requires that X and Y recombine, but they share only a short region of homology called the pseudoautosomal region (PAR) (Raudsepp et al., 2012). In mice, the <1 Mb-long PAR is too small to acquire a DSB in every meiosis if it behaved like a typical genomic segment (genome average of 1 DSB per ∼10 Mb), so the PAR is exceptionally hot for DSB formation (Kauppi et al., 2011; Smagulova et al., 2011; Brick et al., 2012; Lange et al., 2016). Similar to the small chromosomes in yeast, preferential DSB formation in the PAR relies on recruitment of high levels of REC114, MEI4, and IHO1 (the mouse Mer2 ortholog) dependent on intrinsic sequences embedded in the PAR (Acquaviva et al., manuscript in preparation).

It is thus a general principle that the recombination landscape can be shaped profoundly by the intersection between karyotype and the demands of meiotic chromosome segregation, particularly the requirement for at least one DSB per chromosome pair. Budding yeast, with its multilayered chromosome size-dependent control of DSB protein distribution, provides a paradigm for how cells solve the challenge of ensuring recombination on every chromosome, no matter how small.

## Materials and Methods

### Yeast strain construction

#### Myc-epitope tagging of Rec114 and Mer2 in S. cerevisiae and S. mikatae

To perform ChIP-seq experiments in *S. cerevisiae*, Rec114 or Mer2 were C-terminally tagged with 8 and 5 copies of the Myc-epitope marked with the *hphMX4* cassette and the *URA3* gene, respectively (*REC114-Myc-* and *MER2-Myc*) described in (Henderson et al., 2006; Murakami and Keeney, 2014). Rec114 in *S. mikatae* was also C-terminally tagged with the Myc-epitope by the same construct used to tag Rec114 in *S. cerevisiae*. Epitope tagged Rec114 in *S. mikatae* was checked by western blotting and was functional as the *REC114-Myc* strain showed good spore viability (98%, 16 tetrads).

#### Targeting reciprocal translocation between chr1 and chr4

Reciprocal translocation between chr1 and chr4 was targeted as described in **Figure S5A**. The *TRP1* gene with a 3´ portion of the *K. lactis URA3* gene (*Kl.URA3*) was amplified from plasmid pWJ716 (**Key Resources Table)** using primers TL#1AF and TL#1AR (**Key Resources Table)**, which each contain 50 nt from the terminator region of *SWC3* in the left arm of chr1. The *HIS3* gene with a 5´ portion of *Kl.URA3* was amplified from pWJ1077 (**Key Resources Table**) using primers TL#1BF and TL#1BR (**Key Resources Table**), which contain 50 nt from the terminator region of *SLX5* in the left arm of chr4. Each amplified DNA fragment was transformed into *MAT*a and *MATα* haploid yeast (*ura3, trp1, his3*), respectively. After verifying transformants by PCR and sequencing, *MATa* and *MATα* transformants were mated. Since the two *Kl.URA3* segments share identical sequence (448 bp), homologous recombination between these regions would produce uracil prototrophy along with reciprocal translocation between chr1 and chr4. We sporulated the diploid and screened for Ura+ haploids by spreading spores on SC-ura plates. A Ura+ haploid was verified by pulsed-field gel electrophoresis (PFGE) followed by Southern blotting using probes hybridizing to both ends of chr1 and chr4 generated by primers listed in **Key Resources Table** (see **Figure S5B** for an example). We confirmed that native chr1 and chr4 had disappeared and derivative chromosomes der(1) and der(4) of the expected size had appeared. The verified haploid was crossed with a *REC114-myc* haploid which retains native chr1 and chr4 to isolate both mating types with der(1), der(4) and *REC114-myc.* These haploids were verified by Southern blotting again and mated to obtain a diploid with the homozygous translocation in the *REC114-myc* background.

#### Targeting reciprocal translocation between chr8 and chr9

Reciprocal translocation between chr8 and chr9 was targeted by CRISPR/Cas9 as described in **Figure S5A**. Two guide RNA sequences were cloned to pCRCT (*URA3*, iCas9, 2 micron ori, **Key Resources Table**) (Bao et al., 2015) to target cleavages in the downstream regions of YHL012w (80468: chr8, left arm) and *URM1* (342918, chr9, left arm). The plasmid was cotransformed with 100-bp recombination donor fragments that have translocated sequences into a *MATα REC114-myc* haploid. Ura+ transformants were first screened on SC-ura plates and then checked for translocation by PCR with primer pairs flanking the two junctions. Positive transformants were mated with a wild type *MATa* haploid. The resulting diploid turned out to be homozygous for the translocated chromosomes probably because of recombination induced by Cas9 cleavages using der(8) and der(9) as template. The sizes of the translocated chromosomes in the above haploids and diploids were confirmed by PFGE followed by Southern blotting using probes hybridizing to both ends of chr8 and chr9 generated by primers listed in **Key Resources Table** (**Figure S5B**). The diploids that had lost the plasmid were selected on 5-FOA plates and subjected to sporulation followed by tetrad dissection to isolate *MATa* and *MATα* haploids with der(8), der(9), and *REC114-myc*. These haploids were mated and the resulting diploid was used for further experiments.

#### Axis mutants

The *red1, hop1*, and *mek1* deletions were made by replacing the respective coding sequences with the hygromycin B drug resistance cassette (*hphMX4*) amplified from plasmid pMJ696 (identical to pAG32 in (Goldstein and McCusker, 1999). Yeasts were transformed using standard lithium acetate methods. Gene disruption was verified by PCR and Southern blotting. The *SPO11-Flag* construct (*SPO11*-*6His-3FLAG-loxPkanMX-loxP*) was provided by Kunihiro Ohta, Univ. Tokyo (Kugou et al., 2009). All axis (*hop1, red1, rec8, hop1 red1,* and *hop1 red1 rec8*) and *zip3* mutants in the *REC114-Myc* background were created by multiple crossing followed by tetrad dissection.

#### Yeast material and growth conditions

Studies were performed using *S. cerevisiae* SK1 and *S. mikatae* IFO1815 strain backgrounds; strains are listed in in the **Key Resources Table**. Synchronous meiotic cultures were with the SPS pre-growth method described in (Murakami et al., 2009). Saturated overnight cultures in 4 ml YPD (1% yeast extract, 2% peptone, 2% glucose) were used to inoculate 25 ml of SPS (0.5% yeast extract, 1% peptone, 0.67% yeast nitrogen base without amino acids, 1% potassium acetate, 0.05 M potassium biphthalate, pH 5.5, 0.002% antifoam 289 (Sigma)) to a density of 5 × 10_^6^_ cells/ml and cultured at 30°C at 250 rpm for 7 h. Cells were then inoculated into an appropriate volume (900 ml for ChIP-seq experiments with 12 time points or 300 ml for experiments with 3 time points) of fresh SPS at a density of 3 × 10_^5^_ cells/ml and cultured at 30°C at 250 rpm for 12–16 h until the density reached 3–4 × 10_^7^_ cells/ml. Cells were collected by filtration, washed with water, then resuspended at 4 × 10_^7^_ cells/ml in appropriate volume (610 ml for 12 time points or 200 ml for 3 time points) of SPM (2% potassium acetate, 0.001% polypropylene glycol) supplemented with 0.32% amino acid complementation medium (1.5% lysine, 2% histidine, 2% arginine, 1% leucine, 0.2% uracyl, 1% tryptophan). Cultures were shaken at 250 rpm at 30°C and 50 ml samples for ChIP-seq were collected at desired times after transfer to SPM. For 12-time-point cultures, we collected samples as follows: 0, 2, 2.5, 3, 3.5, 4, 4.5, 5, 5.5, 6, 7, 8 h for Rec114 ChIP in *TOF1* background; 0, 1, 1.5, 2, 2.5, 3, 3.5, 4, 4.5, 5, 6, 7 h for Rec114 ChIP in *tof1* background; 0, 0.5, 1, 1.5, 2, 2.5, 3, 3.5, 4, 4.5, 5, 6 h for Mer2 ChIP. For all 3-time-point cultures, cells were collected at 2, 4 and 6 h.

For the Spo11-oligo mapping experiments, synchronous meiotic cultures of *S. cerevisiae* SK1 were prepared as described in (Neale and Keeney, 2009). Briefly, cells from a saturated overnight YPD culture were used to inoculate a 14-h pre-sporulation culture in YPA (1% yeast extract, 2% peptone, 1% potassium acetate) supplemented with 0.001% antifoam 204 and grown at 30°C (starting cell density (OD_600_) of 0.2). Cells were harvested, resuspended in 2% potassium acetate, 0.2 × supplements (2 µg/ml adenine, 2 µg/ml histidine, 6 µg/ml leucine, 2 µg/ml tryptophan, 2 µg/ml uracil), 0.001% antifoam 204 at OD_600_ = 6.0, and incubated in a 30°C shaker to induce sporulation.

To assess culture synchrony, meiotic division profiles were obtained by collecting aliquots at various times from synchronous meiotic cultures, fixing in 50% (v/v) ethanol, and staining with 0.05 µg/ml 4′, 6-diamidino-2phenylindole (DAPI). Mono-, bi- and tetranucleate cells were scored by fluorescence microscopy (data not shown).

### Chromatin immunoprecipitation for Mer2-Myc and Rec114-Myc

We performed ChIP experiments as described in (Murakami and Keeney, 2014), with modifications in cell disruption and chromatin fragmentation. Cells were disrupted by vigorous shaking at 6.5 m/s for 1 min × 10 times in a FastPrep24 (MP Biomedicals). Chromatin in the whole cell extracts (WCE) was sheared by sonication with “M” intensity, 30 sec ON/ 30 sec OFF for 15 min × 3 times in a Bioruptor Sonication System UCD200 (Diagenode) in 15 ml polystyrene conical tubes. Insoluble fraction (cell debris) was removed by centrifugation at 21,130 *g,* 5 min, 4°C. WCE was further sonicated with the same conditions 3–5 times to yield average DNA size less than 500 bp.

For qPCR, we used eight and ten primer pairs for *S. cerevisiae* 12-timepoint and 3-time-points datasets, respectively. For *S. mikatae* we used ten primer sets. All primer sets are listed in **Key Resources Table**. qPCR was performed using the LightCycler^®^ 480 SYBR Green I Master (Roche) according to manufacturer recommendations. All measurements of ChIP samples were expressed relative to the standard (dilution series of corresponding input samples).

### Spo11-oligo mapping

For Spo11-oligo mapping, ≥ 600 ml sporulation culture volume was harvested 4 h after transfer to sporulation media. Because of the severe DSB defect in *red1* and *hop1*, Spo11 oligos from multiple (4–5) cultures of independent colonies were pooled to generate each Spo11-oligo map. The wild-type Spo11-oligo map in **Figure 7C** was from (Mohibullah and Keeney, 2017).

Spo11-oligo mapping in *red1* and *hop1* mutants was performed essentially as described in (Lam and Keeney, 2015), with modifications to purify enough Spo11 oligos from *red1* and *hop1* strains. For example, Spo11 oligos from independent cultures were pooled after eluting from the first immunoprecipitation and at the last step of oligo purification (after proteinase K treatment and ethanol precipitation of the oligos). When pooling Spo11-oligo complexes from five cultures after the first immunoprecipitation step, the total volume of the second immunoprecipitation was increased to 4 ml, and 500 µl of Dynabeads Protein G slurry were pre-bound to 100 µl of 1 mg/ml anti-Flag antibody (as opposed to 125 µl of Dynabeads Protein G slurry pre-bound to 25 µl 1 mg/ml anti-Flag antibody, and a 2^nd^ IP volume of 800 µl). Purified Spo11 oligos were quantified and used for library preparation as described (Thacker et al., 2014).

Sequencing (Illumina HiSeq 2500, 2 × 75 bp paired-end reads) was performed in the MSKCC Integrated Genomics Operation. Clipping of library adapters and mapping of reads was performed by the Bioinformatics Core Facility (MSKCC) using a custom pipeline as described (Pan et al., 2011; Thacker et al., 2014; Lam and Keeney, 2015; Zhu and Keeney, 2015; Mohibullah and Keeney, 2017). Reads were mapped to the sacCer2 genome assembly of type strain S288C from SGD (*Saccharomyces* Genome Database).

### ChIP-seq data processing: scaling and masking

ChIP-seq experiments were performed as described in (Murakami and Keeney, 2014). DNA from ChIP and input samples (same samples as used for ChIP-qPCR) were further sheared by sonication to an average fragment sizes of ∼300 bp. These were sequenced (50 bp paired-end) on the HiSeq platform (Illumina). Reads were mapped to the SacCer2 genome assembly and *S. mikatae* genome assembly from (Scannell et al., 2011) using BWA (0.7) MEM to generate coverage maps for each time point from each strain. Each ChIP coverage map was divided by the corresponding input map for normalization. Then, to scale the ChIP-seq coverage relative to absolute ChIP efficiency, we calculated the total coverage within ± 1 kb of the center of each qPCR amplicon, plotted these as a function of the corresponding qPCR ChIP efficiency, and calculated regression lines by least squares. The resulting regression line for each time point was then used to scale the ChIP-seq coverage maps.

To remove regions with spurious mapping, we previously defined “mask regions” where the coverage from the 0 h sample of the wild-type *ARS+* strain was out of a fixed range (>1.5 SD from mean coverage) with further extension by 1 kb on other side. These regions were censored in all input and ChIP coverage maps from *S. cerevisiae*. Mask regions for *S. mikatae* were defined similarly where the coverage from 2 h input sample exceeded a fixed range (mean coverage ± 4 SD, calculated between 50–150 kb region of chr1). After the same extension, these regions were censored from *S. mikatae* coverage maps.

### Replication index generated by ChIP input coverage maps

All masked coverage maps from input samples were binned using 5 kb windows and normalized to genomic mean coverage. For 12-time-point datasets, coverage from an “S-phase time point” (1.5 h and 2.5 h for Rec114 ChIP *ARS+ tof1Δ* and Rec114 ChIP *arsΔ tof1Δ,* respectively; 2 h for the rest) was divided by the corresponding “G1-phase time point” (0-h sample) to generate a “relative coverage” map. For 3-time-point datasets, the 2-h time point map was divided by the 0 h map from the Rec114 ChIP *ARS+* dataset to generate relative coverage. For the *S. mikatae* dataset, the mean normalized 2 h map was used as relative coverage. We defined the “replication index” as –log_2_(relative coverage). Outliers were removed from each dataset, defined as the replication index value exceeding a fixed range (mean ± 4 SD).

### Estimating association and dissociation times by sequential curve fitting

The method to measure association time is described in (Murakami and Keeney, 2014). The scaled and masked ChIP-seq coverage maps from two Rec114 ChIP-seq and Mer2 ChIP-seq data sets were smoothed using a 2010-bp Parzen (triangular) sliding window. Using the smoothed, scaled coverage map at 3.5 h time points, a total of 1477 (Rec114 ChIP *ARS+*), 1545 (Rec114 ChIP *arsΔ*) and 1550 (Mer2 ChIP) peaks were called using as a threshold 0.5× each chromosome’s mean coverage. A ChIP temporal profile at each peak position was assembled by collecting the ChIP signals from the smoothed, scaled coverage map for each time point.

To define the empirical maximum position in the ChIP profile (t_max_), Gaussian curves were fitted to ChIP signals plotted as a function of time. To create positive skew in the regression curves, times (t, in hours) were log-transformed [t′ = ln(t+1)]. We employed an equation that is a modification of the Gaussian probability density function:

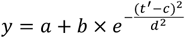

where *y* is ChIP signal, *a* is the background, *b* is the peak height, *c* is the peak position, and *d* is the equivalent of standard deviation. We set the background parameter (*a*) to the ChIP signal at 0 h, then fitted the equation to the data points by least squares to estimate the other parameters (*b, c* and *d*) using the “nls” function in R. The estimated parameter (c) was transformed back to h in meiosis (t_max_ = e^c^-1).

Next, to estimate the association time of DSB protein, we used this peak to fit a saturating exponential growth (logistic) curve to just the upward slope of the ChIP temporal profile (data points before t_max_):

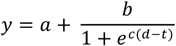

where *y* is ChIP signal, *a* is the background, *b* is the maximum value, *c* is a shaping factor and *d* is the inflection point of the logistic function, respectively. We set the background and the maximum value parameters (*a* and *b*) to the ChIP signal at 0 h and the previously estimated peak height value (parameter (*b*) from skewed Gaussian fitting, *b_Gauss_*), respectively, and then fitted the equation to the data points to estimate the other parameters (*c* and *d*) using the “nls” function in R. We used *d* as t_association_ where the logistic curve reaches 50% of maximum.

We also estimated the dissociation time of DSB protein by fitting a logistic curve to the downward slope of the ChIP temporal profile (data points after t_max_):

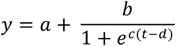

where *y* is ChIP signal, *a* is the background, *b* is the maximum value, *c* is a shaping factor and *d* is the inflection point of the logistic function, respectively. We used the “nls” function to estimate parameters (*c* and *d*) and used *d* as t_dissociation_ where the logistic curve reaches 50% of maximum.

To evaluate the fitting quality for the kinetic profile at each peak, absolute distances between the data points and the fitted Gaussian curve (residuals) were summed and divided by the peak height (parameter *bGauss*) from the fitted curve (normalized-total residuals, *r*_Gauss_). Total residuals from two logistic fittings were divided by *b_Gauss_*, and the sum of these was defined as *r*_logistic_. We excluded poorly fitted peaks with normalized residuals exceeding 1.2 for Rec114 ChIP *arsΔ* and Mer2 ChIP datasets. We used less stringent criteria (filtering out peaks with *r*_Gauss_ > 1.6 or *r*_logistic_ > 1.5) for Rec114 ChIP *ARS+* dataset because overall quality of fittings was less good compared to the other two datasets. After these filtering steps, totals of 998 (Rec114 ChIP *ARS+*), 1081 (Rec114 ChIP *arsΔ*), and 1490 (Mer2 ChIP) peaks were processed for further analyses.

For the Rec114 association time in the two *tof1* datasets, we used previously estimated values for 957 (*tof1 ARS+*) and 2020 (*tof1 arsΔ*) peaks that passed filtering (Murakami and Keeney, 2014).

### Estimating centromere and telomere effects on DSB protein association time and ChIP density at 2 h

Association timing and ChIP density (2 h) data from the two Rec114 and one Mer2 ChIP time courses were combined as follows. To capture the intra-chromosomal features separate from inter-chromosomal differences, we averaged each dataset in 20-kb bins, standardized the values for each chromosome to a mean of 0 and variance of 1, and standardized again within a dataset before pooling all datasets together. The first standardization minimizes differences between chromosomes and the second standardization minimizes differences between datasets. The pooled association time or ChIP density (2 h) data were plotted as a function of distance from centromere or telomere. We fitted an exponential decay model to these data points:

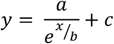

where *y* is association time or ChIP density at 2 h, *x* is distance from centromere or telomere, *a* is the initial value of the centromere or telomere effect, *b* is a shaping factor and *c* is an intercept, respectively. We used the “nls” function in R to estimate parameters (*a, b* and *c*) and used the parameter (*b*) to present the half distance where the initial value decays to half as the following:

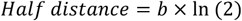

### Multiple linear regression analysis

We performed multiple linear regression analyses using replication index and the centromere and telomere effects as explanatory variables and either association time or ChIP density at 2 h as response variables. To perform multiple regression analyses, replication index and association time or ChIP density at 2 h were averaged in 20-kb bins. Distances from centromere and telomere at the midpoints of the 20-kb bins were plugged into the centromere and telomere exponential models whose parameters were estimated as described in the preceding section using the pooled, standardized data for association or ChIP density at 2 h. The same centromere and telomere models were used for multiple regression in strains with translocation and in *S. mikatae*. Estimates of the regression coefficients and the standardized regression coefficients (beta) are shown, along with *t* and *P* values based on the standardized coefficients in **Supplementary Table 1**.

### Data and software availability

All sequencing data were deposited at the Gene Expression Omnibus (GEO) with the accession numbers GSE52970 (Rec114 ChIP-seq including *tof1*), GSE84859 (Spo11 oligos in *hop1* and *red1*). Accession numbers pending for Mer2 ChIP-seq and all Rec114 ChIP-seq generated in this study and for Spo11-oligo maps in wild type at 4 and 6 h.

## End Matter

### Author Contributions and Notes

HM performed ChIP-seq, generated translocation strains, and analyzed the data. IL and MvO performed Spo11-oligo mapping. JS performed ChIP-seq under the supervision of HM. HM and SK conceived the project and wrote the paper. SK analyzed data, procured funding, and oversaw the research. HM, SK, and IL edited the manuscript.

The authors declare no competing interests.

## Acknowledgments

We are grateful to Agnès Viale and Neeman Mohibullah of the Memorial Sloan Kettering Cancer Center (MSKCC) Integrated Genomics Operation for DNA sequencing; Nicholas Socci at the MSKCC Bioinformatics Core Facility for mapping ChIP-seq and Spo11-oligo reads; and members of the Keeney laboratory, especially Shintaro Yamada for advice on data analysis and Laurent Acquaviva for sharing unpublished information. We thank Vijayalakshmi Subramanian (NYU), Andreas Hochwagen (NYU), and Franz Klein (Univ. of Vienna) for discussions and sharing unpublished information. We thank Michael Lichten (NCI), Ed Louis (Univ. of Nottingham), Kunihiro Ohta (Tokyo Univ.), and Rodney Rothstein (Columbia Medical Center) for strains or plasmids. IL and MvO were supported in part by National Institutes of Health (NIH) fellowships F31 GM097861 and F32 GM096692, respectively. This work was supported by NIH grants R01 GM058673 and R35 GM118092 to SK. MSKCC core facilities are supported by NCI Cancer Center Support Grant P30 CA008748.

---

**Figure S1.**
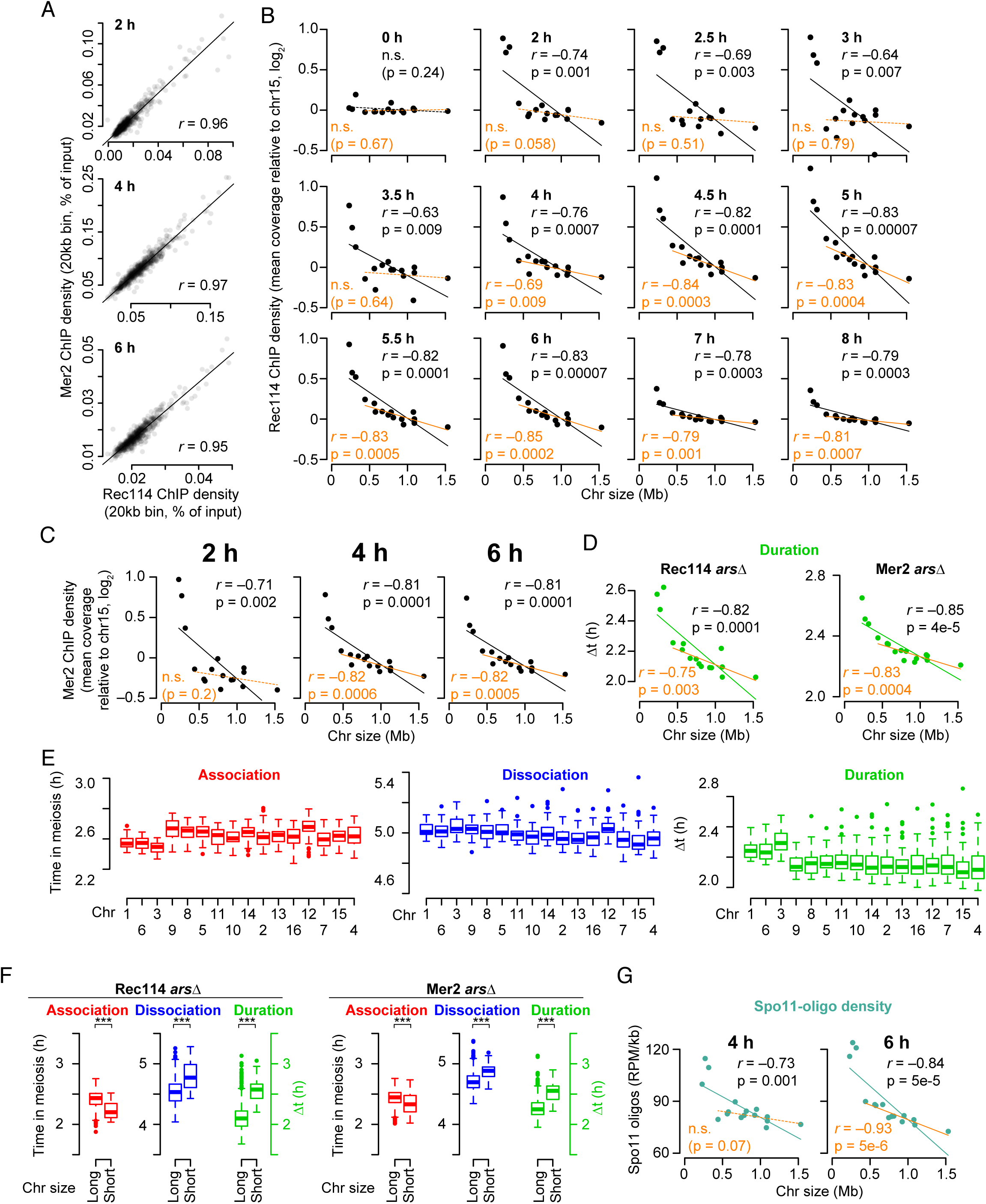
Chromosome size dependence of DSB protein binding, related to Figure 1. (A) Similarity of Rec114 and Mer2 ChIP-seq patterns. Both strains are *arsΔ* background. (B) Full time course of average per-chromosome Rec114 ChIP densities (*ARS+* strain). (C) The size dependence of average per-chromosome Mer2 ChIP density changes over time similar to Rec114 (compare to Figure 1E). (D) Per-chromosome mean Rec114 and Mer2 binding duration in *arsΔ* strains. (E) Association times, dissociation times, and binding duration of Rec114 (*ARS+* strain) at Rec114 peaks, broken down by chromosome. (F) Comparison between the shortest three chromosomes and the other thirteen for DSB protein association, dissociation and duration (remaining two datasets to accompany Figure 1I). (G) Chromosome size dependency of Spo11-oligo distributions as a function of sampling time.

**Figure S2.**
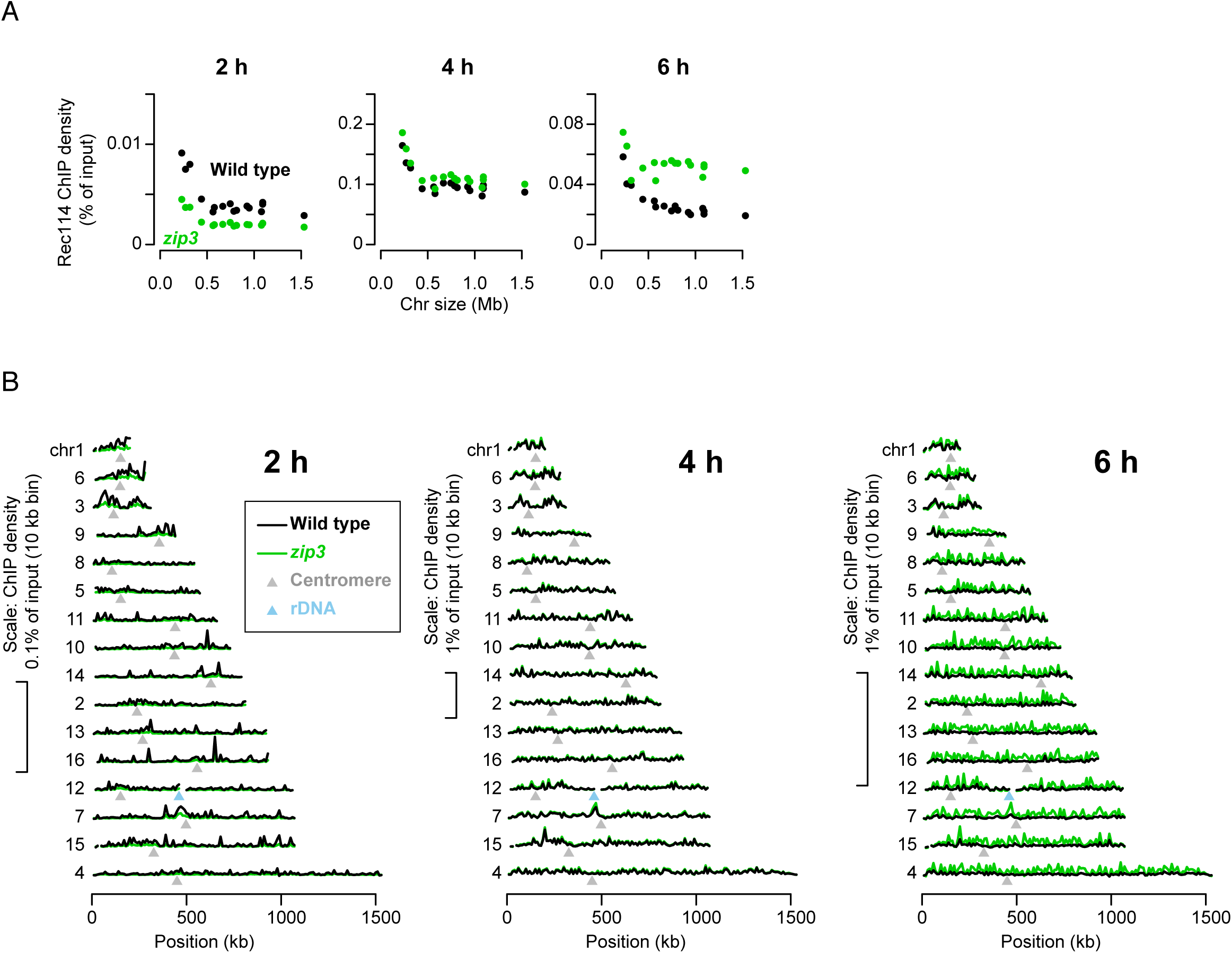
Rec114 chromatin binding in the *zip3* mutant, related to Figure 2. (A) Effects of *zip3* mutation on absolute Rec114 ChIP densities. ChIP-seq coverage was calibrated by qPCR. Chromosomal means are shown. At 2 h, which is very early in the initial accumulation phase of Rec114 on chromatin, the *zip3* mutant has ∼two-fold less Rec114 signal, affecting all chromosomes equivalently. It is possible that this decrease reflects a role for Zip3 early, but since Zip3 is not known to have any function this early in prophase, a more likely explanation is that the *zip3* mutant culture lagged slightly behind the wild-type culture. At 4 h, Rec114 ChIP densities were highly similar between the cultures, again with little or no difference between chromosomes. However, at 6 h, the *zip3* mutant retained substantially more Rec114 signal. At least some of this increase may be attributable to delayed meiotic progression because of inhibition of Ndt80 (Carballo et al., 2013; Thacker et al., 2014), but importantly for purposes of this experiment, the Rec114 levels are disproportionately elevated on the 13 larger chromosomes. (B) Per-chromosome profiles of absolute Rec114 ChIP density.

**Figure S3.**
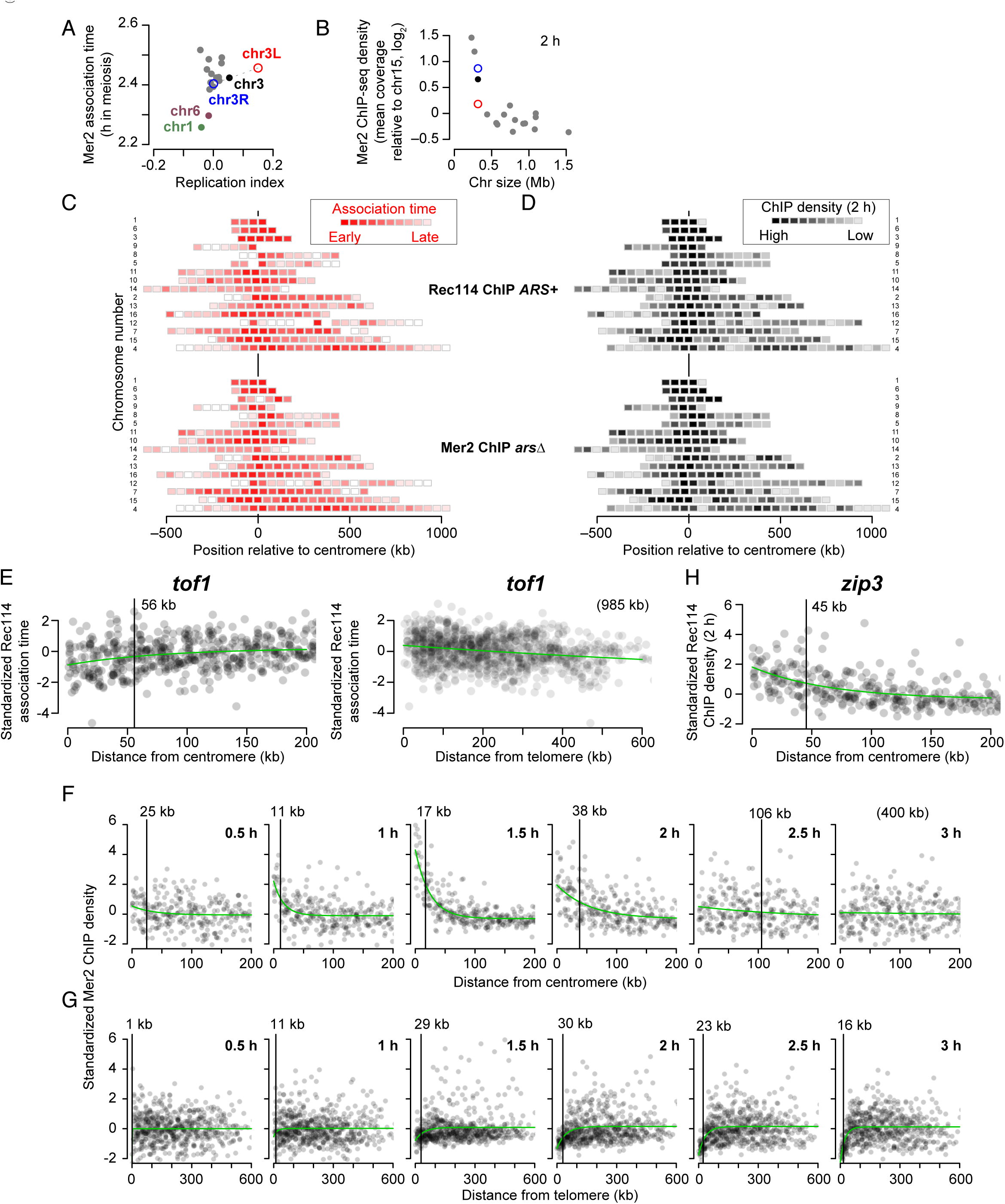
Replication timing and the centromere and telomere effects, related to Figure 3. (A) Comparison of per-chromosome Mer2 association time with replication timing. Compare with Rec114 datasets in Figure 3A. The similarity with Rec114 patterns, including much later Mer2 accumulation on the late-replicating left arm of chr3, suggests that Mer2 binding to chromatin is also coordinated with replication timing. Mer2 is able to bind chromatin in the absence of Rec114 (Henderson et al., 2006; Li et al., 2006; Panizza et al., 2011), but interaction with Rec114 (which is promoted by replication-associated Rec114 phosphorylation (Sasanuma et al., 2008)) might stabilize or otherwise modify the localization of Mer2. (B) Average per-chromosome Mer2 ChIP density at 2 h (normalized to chr15). Compare with Rec114 datasets in Figure 3B. (C,D) Intra-chromosomal distributions of Rec114 and Mer2 association times (C) and ChIP density at 2 h (D) in the indicated strains. Average values for 50-kb bins are presented as described in the legend to Figures 3C and 3D. (E) Centromere (left graph) and telomere (right graph) effects in *tof1* mutants. Rec114 ChIP-seq data from *ARS+ tof1* and *arsΔ tof1* strains were binned, standardized, and pooled as described in the legend to Figure 3E. The centromere and telomere effects are still apparent in the *tof1* mutants, but appear substantially weaker (particularly the telomere effect). (F, G) Detailed time courses of Mer2 binding to chromatin near centromeres (F) and telomeres (G). Mer2 ChIP density was binned and standardized as described in the legend to Figure 3E.

**Figure S4.**
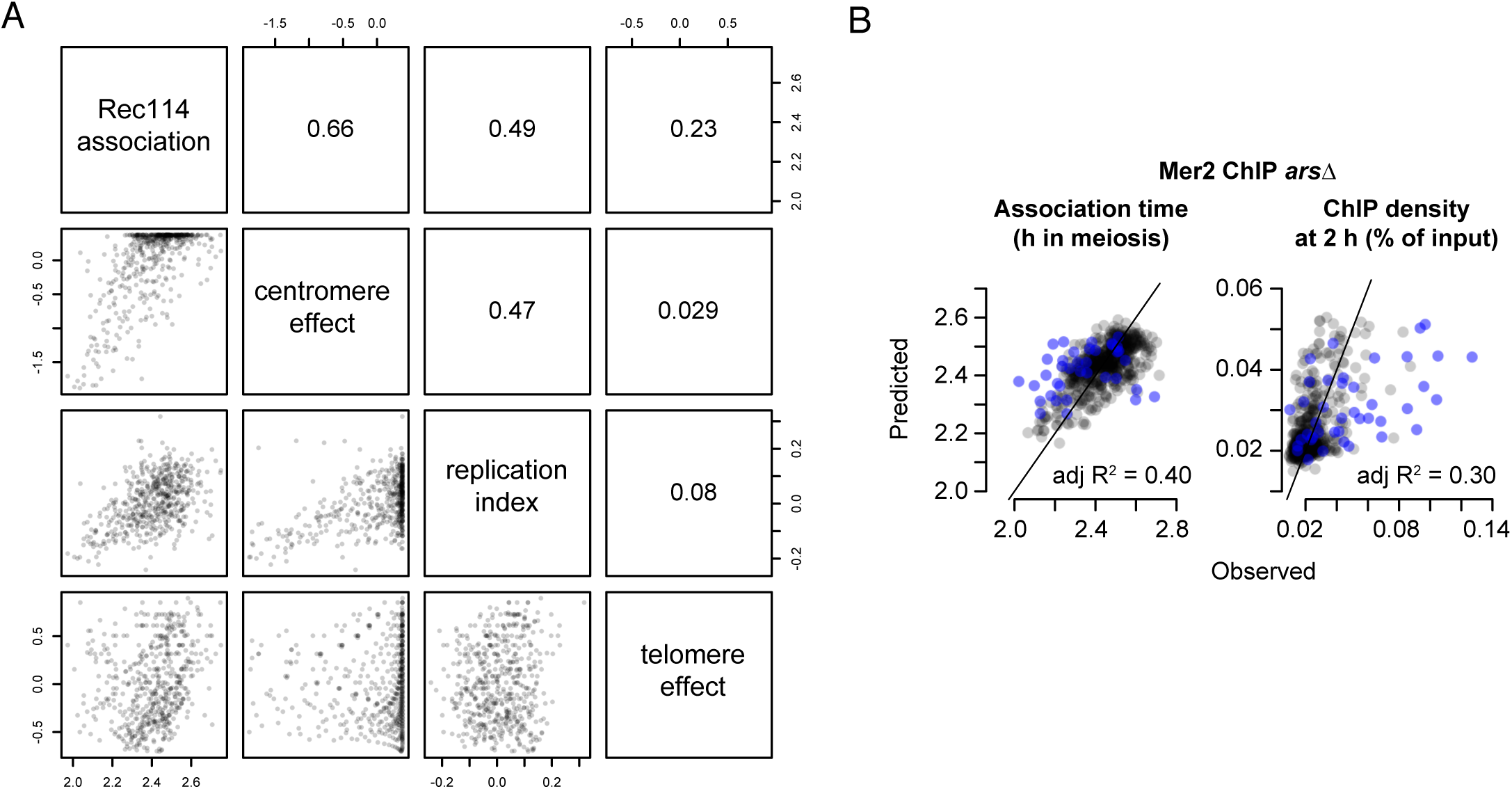
Multiple regression model integrating replication timing and the centromere and telomere effects, related to Figure 4. (A) Pairwise correlations between Rec114 association time, replication index, and centromere and telomere effects. Numbers in boxes indicate Pearson’s correlation coefficients. Data are from the *ARS+* strain. (B) Observed Mer2 association times (left) or ChIP density at 2 h (right) vs. values predicted from a three-factor multiple regression model. Mer2 ChIP data gave similar results as Rec114 (Figures 4A and 4D). Each point represents a 20-kb bin. Points for bins on short chromosomes are blue.

**Figure S5.**
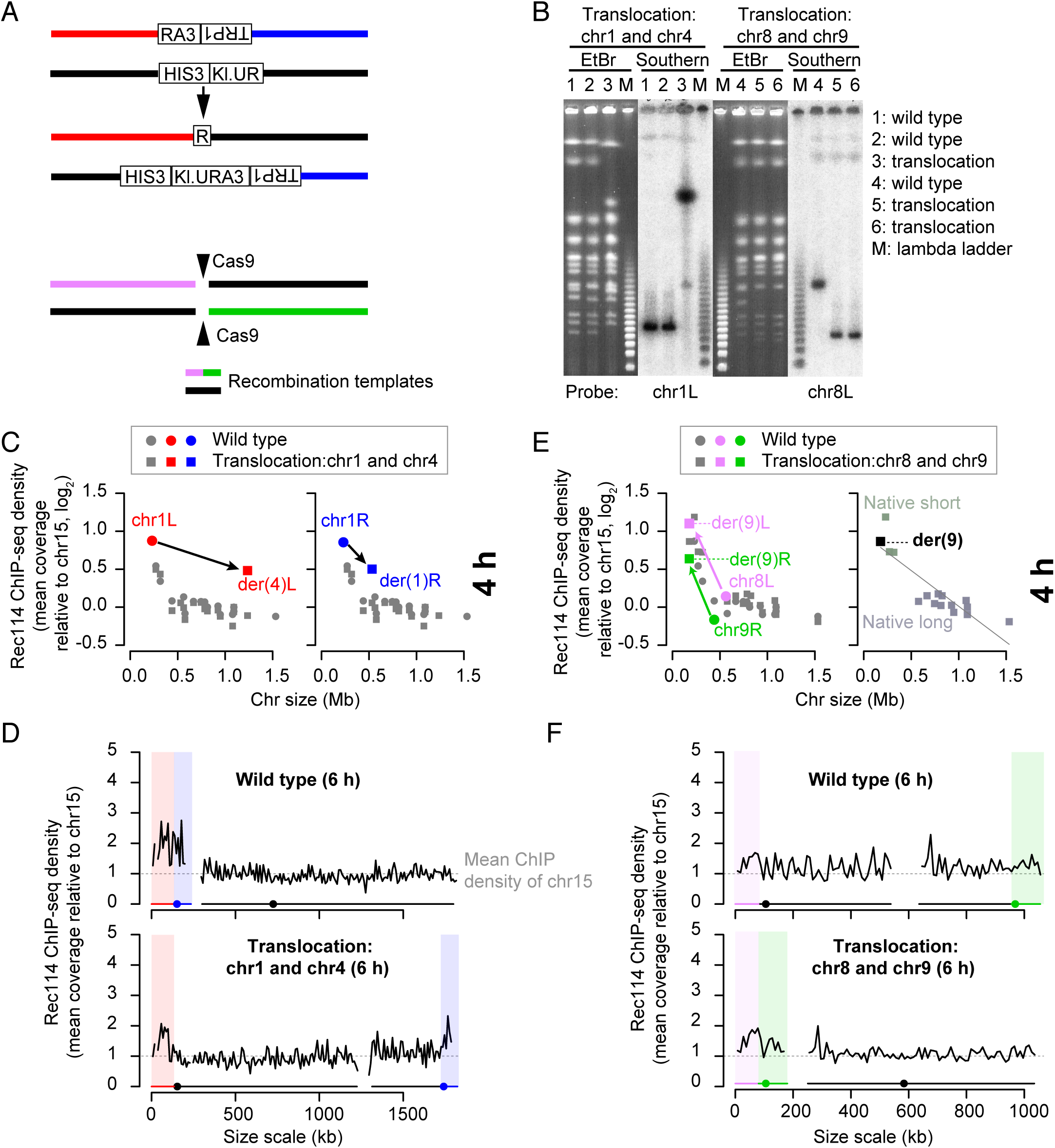
Effects of targeted translocations on per-chromosome Rec114 binding, related to Figure 5. (A) Strategies to target reciprocal translocations. Top: to generate the translocation between chr1 and chr4, part of the 3′ end of the *URA3* gene from *Kluyveromyces lactis* (“*RA3*”) was integrated on chr1 along with the *TRP1* gene. Separately, part of the 5′ end of *K. lactis URA3* (“*Kl.UR”)* was integrated on chr4 along with the *HIS3* gene. The two parts of *K. lactis URA3* partially overlap, so their shared region of homology allows reciprocal recombination between them to result in formation of a functional *URA3* gene. Bottom: to generate the translocation between chr8 and chr9, we introduced a plasmid expressing Cas9 and two guide RNAs that target cleavage within chr8 and chr9, respectively. This plasmid was introduced by co-transforming it along with 100 bp long recombination templates matching the desired reciprocal recombination products. (B) Confirmation of targeted translocations. High-molecular weight DNA was prepared from control and translocation strains and separated on pulsed-field gels and stained with ethidium bromide (EtBr). The translocations were then verified by Southern blotting using probes against the right and left ends of both chromosomes involved in the translocation. Representative results using the chr1L and chr8L probes are presented. (C) Chr1-derived sequences retain high-level Rec114 binding when in a large-chromosome context. Average per-chromosome Rec114 ChIP densities normalized to chr15 are shown at 4 h (the intermediate time point for the samples shown in Figure 5B). (D) Rec114 profiles for wild-type and translocated chromosomes at 6 h. ChIP-seq data were normalized relative to chr15 and smoothed with a 10-kb sliding window. (E,F) An artificially short chromosome fails to acquire a boost in Rec114 binding. Per-chromosome average Rec114 ChIP densities (F) and Rec114 profiles (G) are shown at 4 h (the intermediate time point for the samples shown in Figure 5F).

**Figure S6.**
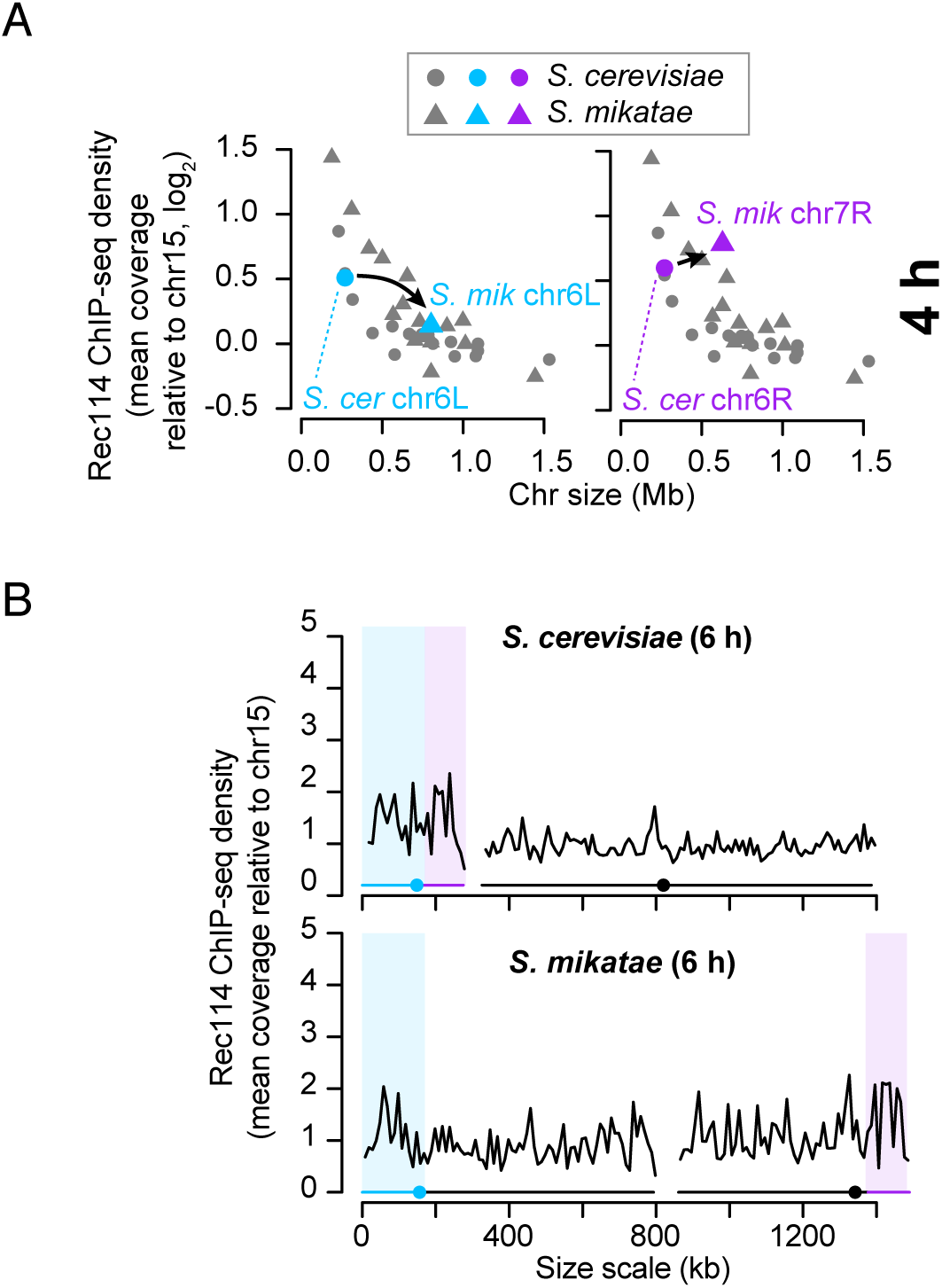
High-level Rec114 binding to chr6-derived sequences is not retained in *S. mikatae*, related to Figure 6. Rec114 ChIP density at 4 h (A) and Rec114 profiles (B) are shown for additional time points, as in Figures 5B and 5C, respectively.

**Figure S7.**
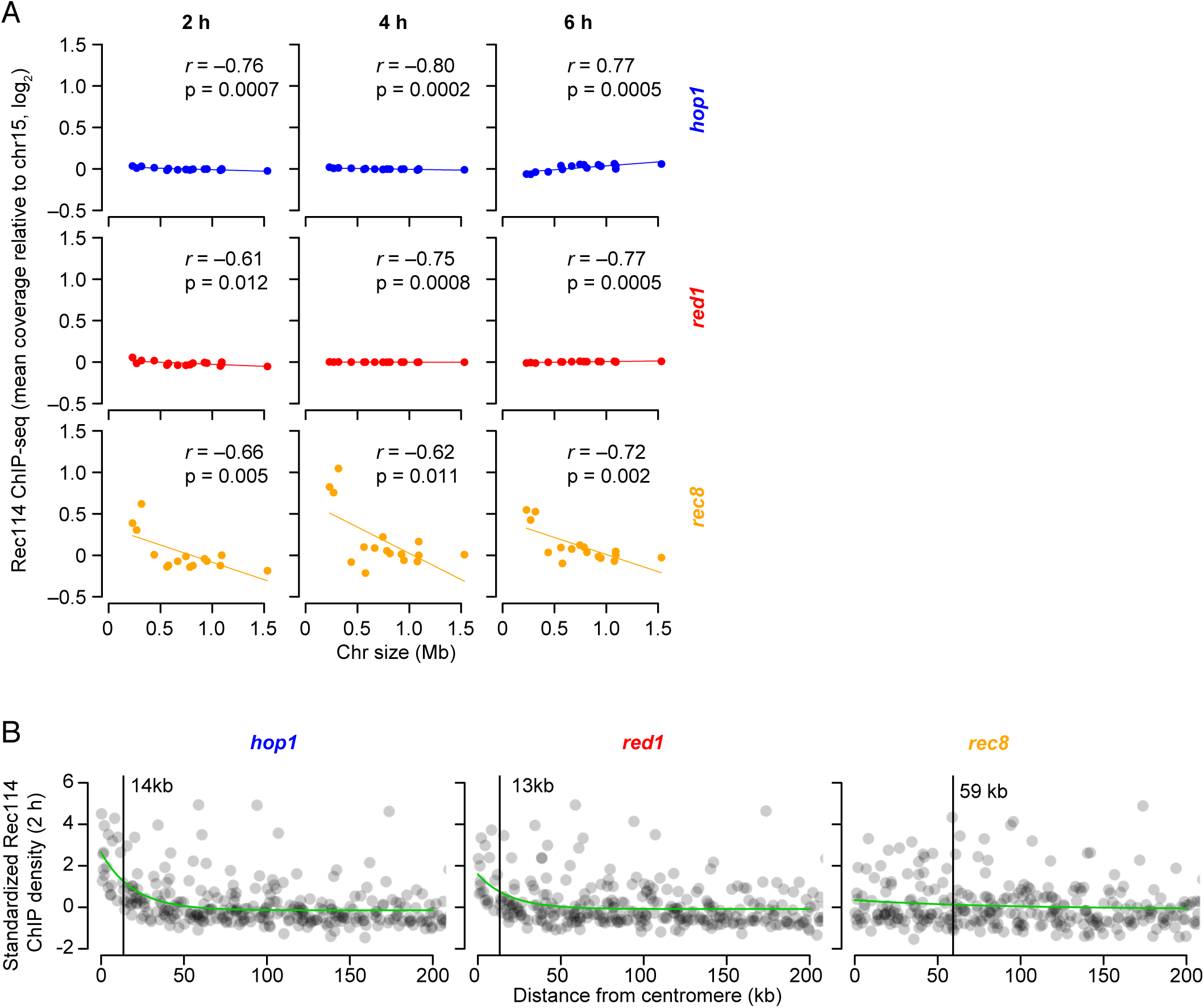
Effect of chromosomes axis proteins on Rec114 chromatin binding patterns, related to Figure 7. (A) Chromosome size dependence of Rec114 binding to chromosomes is lost in the absence of Hop1 or Red1, but not Rec8. Results from single mutants are presented as in Figure 7B. Note that, although correlations with chromosome size remain statistically significant in both the *hop1* and *red1* mutants, their slopes are negligible compared to wild type (Figure 1E). (B) The centromere effect is retained (albeit spreading less far) in *hop1* and *red1* single mutants. Rec114 ChIP data at 2 h were binned and standardized as described for Figure 3E.

**Supplementary Table 1.**
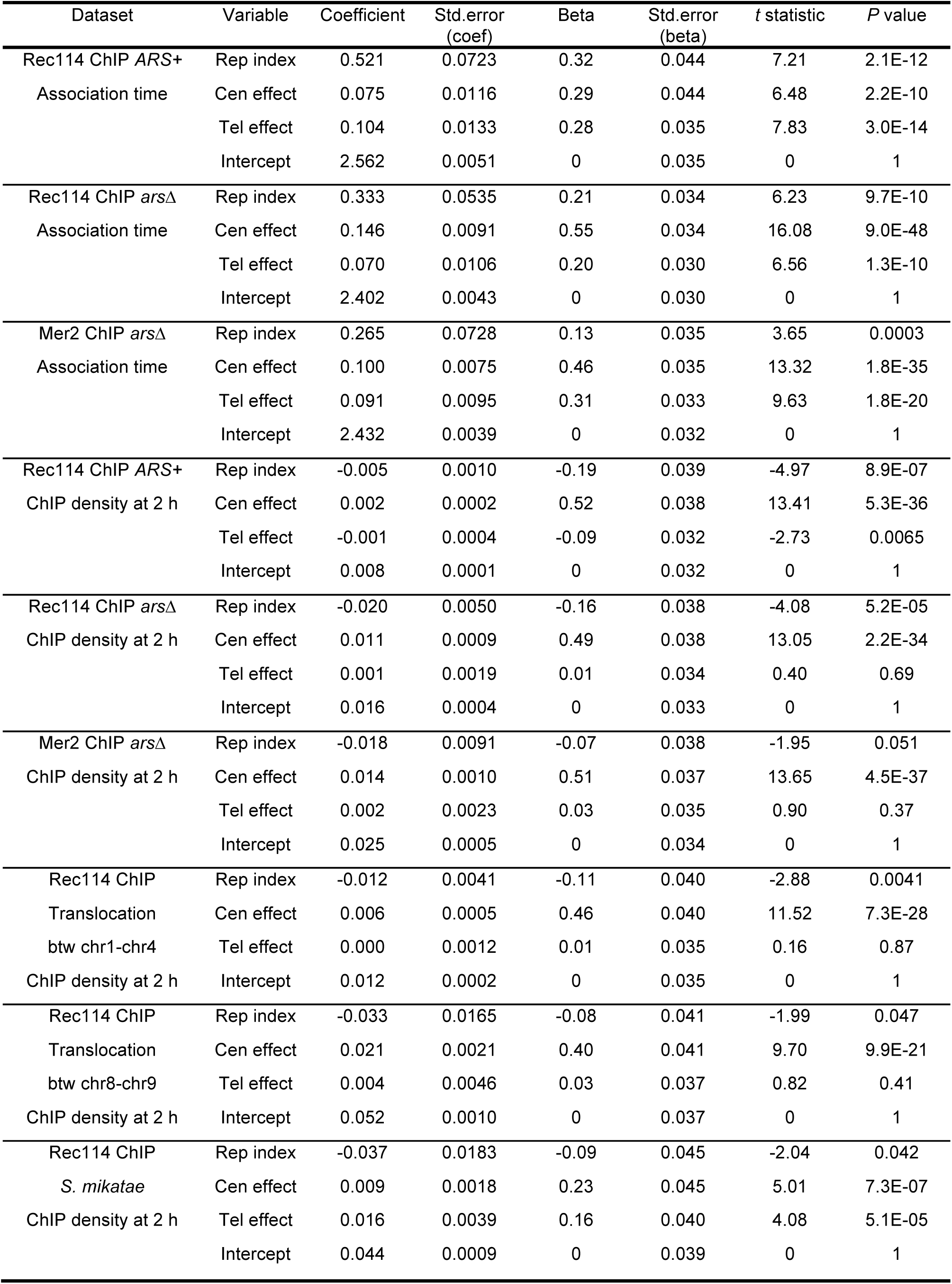
Multiple regression results. Hajime Murakami, Isabel Lam, Jacquelyn Song, Megan van Overbeek & Scott Keeney Multilayered mechanisms ensure that short chromosomes recombine in meiosis

## KEY RESOURCES TABLE

Hajime Murakami, Isabel Lam, Jacquelyn Song, Megan van Overbeek & Scott Keeney Multilayered mechanisms ensure that short chromosomes recombine in meiosis

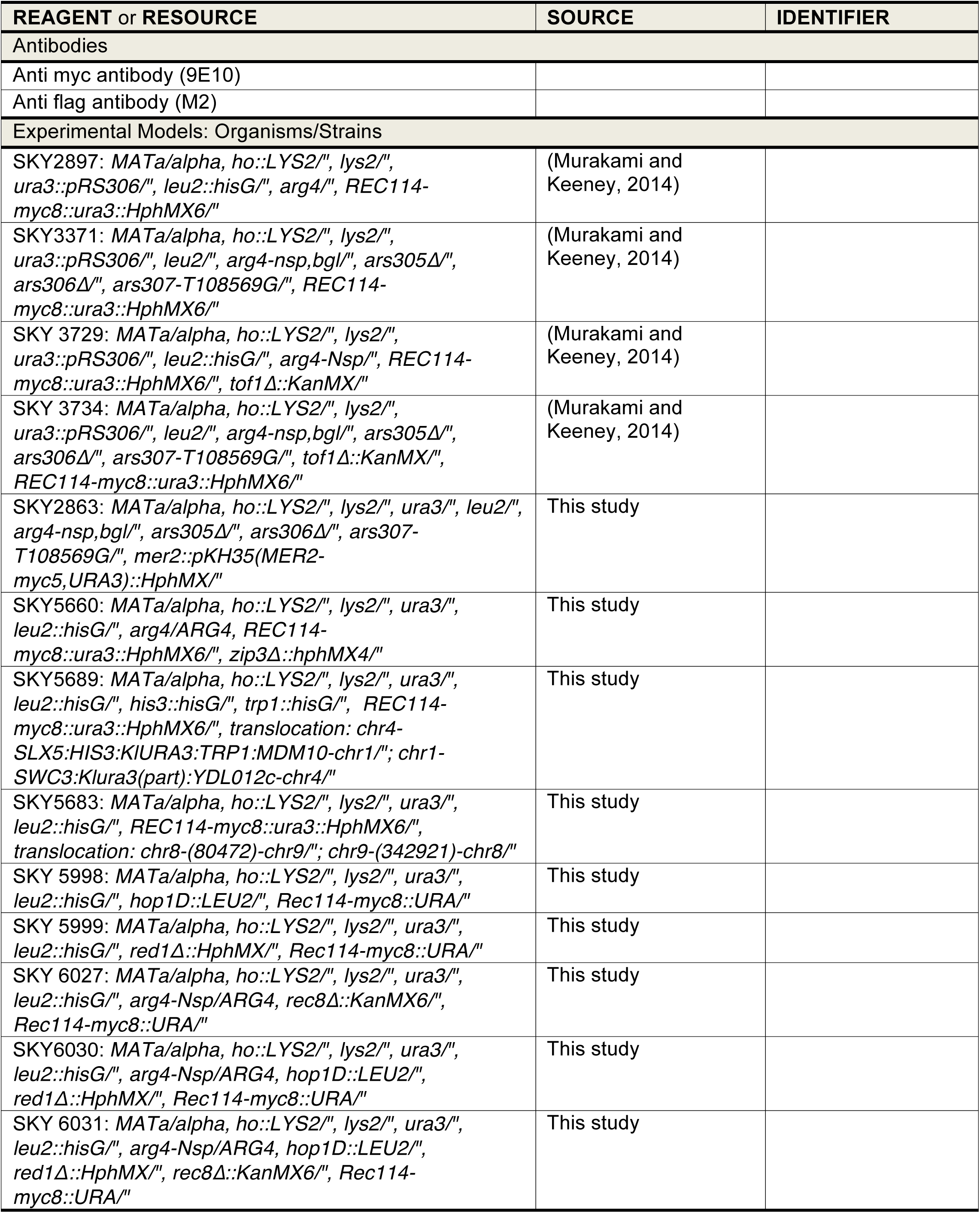
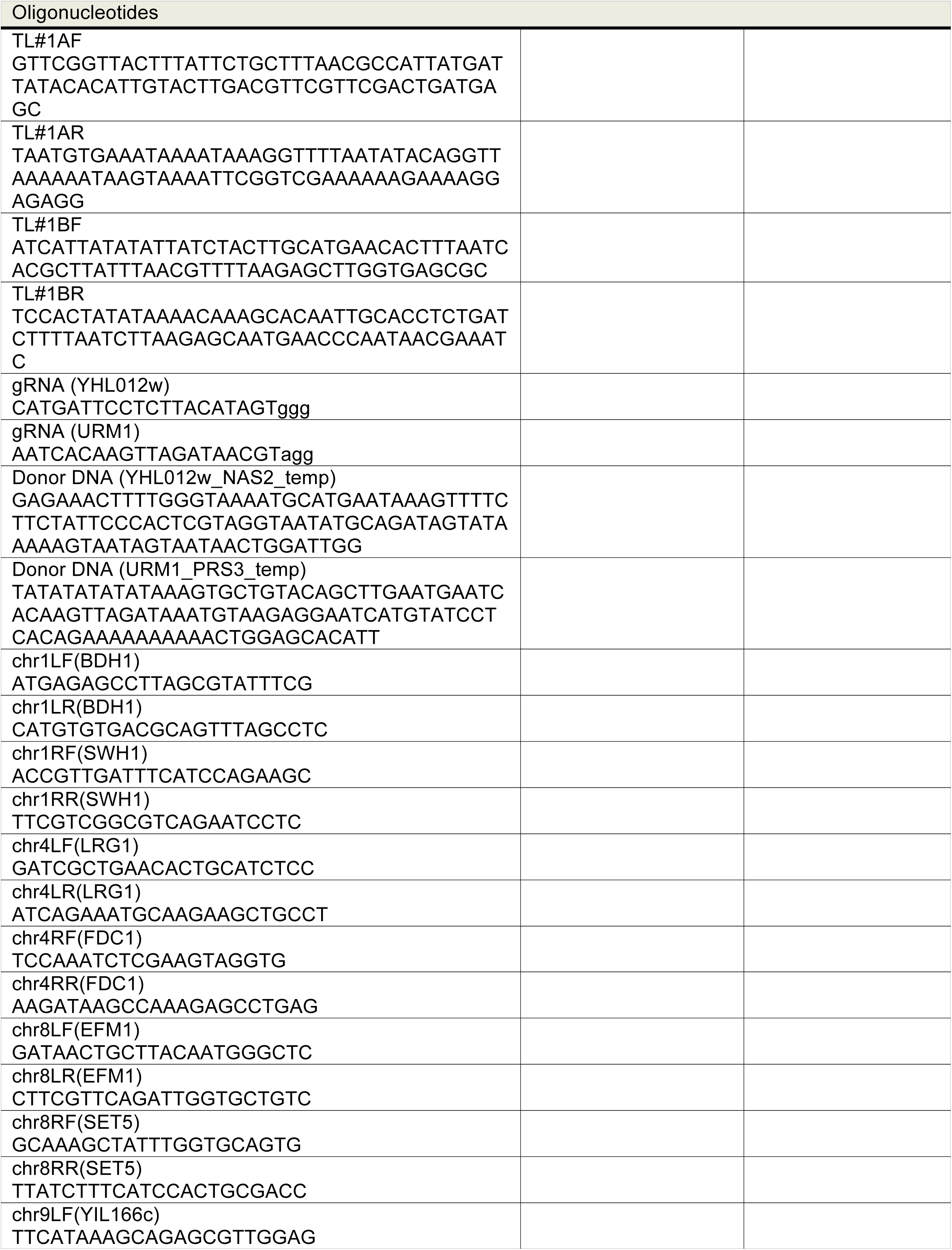
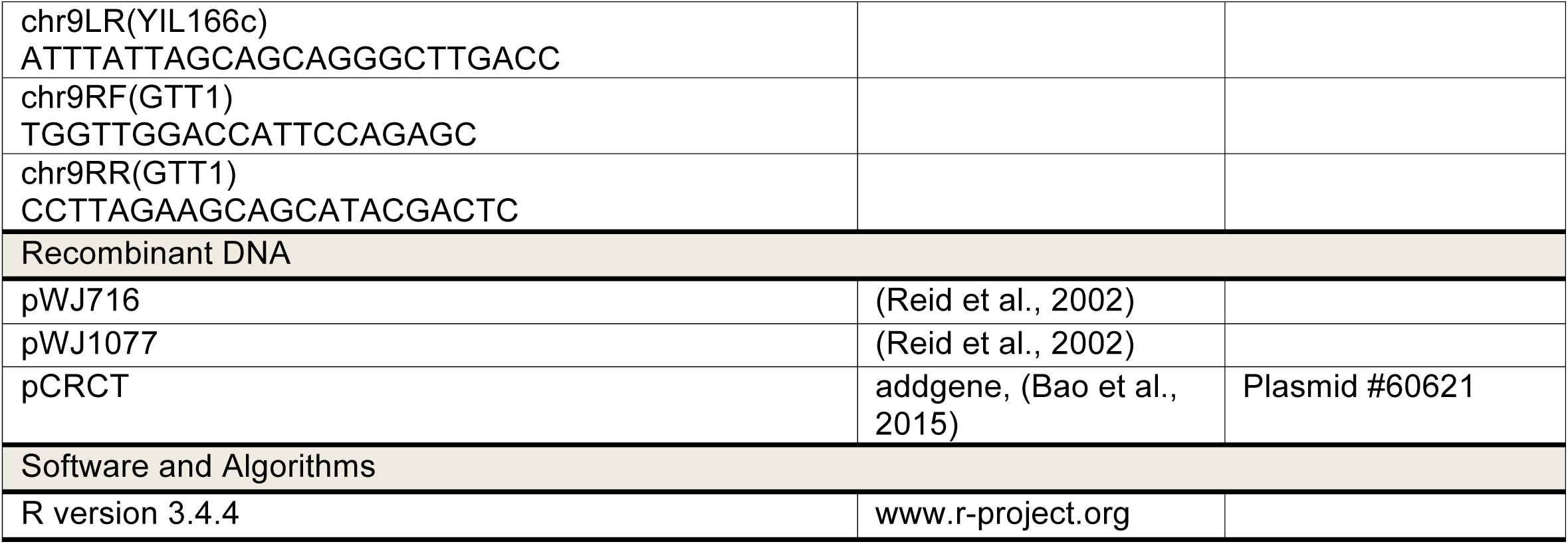

## References

Arora, C., Kee, K., Maleki, S., and Keeney, S. (2004). Antiviral protein Ski8 is a direct partner of Spo11 in meiotic DNA break formation, independent of its cytoplasmic role in RNA metabolism. Mol Cell 13, 549–559.

Bao, Z., Xiao, H., Liang, J., Zhang, L., Xiong, X., Sun, N., Si, T., and Zhao, H. (2015). Homology-integrated CRISPR-Cas (HI-CRISPR) system for one-step multigene disruption in Saccharomyces cerevisiae. ACS Synth Biol 4, 585–594.

Baudat, F., and Nicolas, A. (1997). Clustering of meiotic double-strand breaks on yeast chromosome III. Proc Natl Acad Sci U S A 94, 5213– 5218.

Blat, Y., Protacio, R.U., Hunter, N., and Kleckner, N. (2002). Physical and functional interactions among basic chromosome organizational features govern early steps of meiotic chiasma formation. Cell 111, 791– 802.

Blitzblau, H.G., Chan, C.S., Hochwagen, A., and Bell, S.P. (2012). Separation of DNA replication from the assembly of break-competent meiotic chromosomes. PLoS Genet 8, e1002643.

Borde, V., Goldman, A.S., and Lichten, M. (2000). Direct coupling between meiotic DNA replication and recombination initiation. Science 290, 806–809.

Brick, K., Smagulova, F., Khil, P., Camerini-Otero, R.D., and Petukhova, G.V. (2012). Genetic recombination is directed away from functional genomic elements in mice. Nature 485, 642–645.

Carballo, J.A., Panizza, S., Serrentino, M.E., Johnson, A.L., Geymonat, M., Borde, V., Klein, F., and Cha, R.S. (2013). Budding yeast ATM/ATR control meiotic double-strand break (DSB) levels by down-regulating Rec114, an essential component of the DSB-machinery. PLoS Genet 9, e1003545.

Chen, S.Y., Tsubouchi, T., Rockmill, B., Sandler, J.S., Richards, D.R., Vader, G., Hochwagen, A., Roeder, G.S., and Fung, J.C. (2008). Global analysis of the meiotic crossover landscape. Dev Cell 15, 401–415.

Cooper, T.J., Wardell, K., Garcia, V., and Neale, M.J. (2014). Homeostatic regulation of meiotic DSB formation by ATM/ATR. Exp Cell Res 329, 124–131.

Fischer, G., James, S.A., Roberts, I.N., Oliver, S.G., and Louis, E.J. (2000). Chromosomal evolution in *Saccharomyces*. Nature 405, 451–454.

Goldstein, A.L., and McCusker, J.H. (1999). Three new dominant drug resistance cassettes for gene disruption in Saccharomyces cerevisiae. Yeast 15, 1541–1553.

Gray, S., Allison, R.M., Garcia, V., Goldman, A.S., and Neale, M.J. (2013). Positive regulation of meiotic DNA double-strand break formation by activation of the DNA damage checkpoint kinase Mec1(ATR). Open Biol 3, 130019.

Henderson, K.A., Kee, K., Maleki, S., Santini, P.A., and Keeney, S. (2006). Cyclin-dependent kinase directly regulates initiation of meiotic recombination. Cell 125, 1321–1332.

Henderson, K.A., and Keeney, S. (2004). Tying synaptonemal complex initiation to the formation and programmed repair of DNA double-strand breaks. Proc Natl Acad Sci U S A 101, 4519–4524.

Joshi, N., Brown, M.S., Bishop, D.K., and Borner, G.V. (2015). Gradual implementation of the meiotic recombination program via checkpoint pathways controlled by global DSB levels. Mol Cell 57, 797–811.

Kaback, D.B., Guacci, V., Barber, D., and Mahon, J.W. (1992). Chromosome size-dependent control of meiotic recombination. Science 256, 228–232.

Kauppi, L., Barchi, M., Baudat, F., Romanienko, P.J., Keeney, S., and Jasin, M. (2011). Distinct properties of the XY pseudoautosomal region crucial for male meiosis. Science 331, 916–920.

Kauppi, L., Barchi, M., Lange, J., Baudat, F., Jasin, M., and Keeney, S. (2013). Numerical constraints and feedback control of double-strand breaks in mouse meiosis. Genes Dev 27, 873–886.

Keeney, S. (2007). Spo11 and the Formation of DNA Double-Strand Breaks in Meiosis. In Recombination and Meiosis, D.H. Lankenau, ed. (Heidelberg, Germany: Springer-Verlag), pp. 81–123.

Keeney, S., Giroux, C.N., and Kleckner, N. (1997). Meiosis-specific DNA double-strand breaks are catalyzed by Spo11, a member of a widely conserved protein family. Cell 88, 375–384.

Keeney, S., Lange, J., and Mohibullah, N. (2014). Self-organization of meiotic recombination initiation: general principles and molecular pathways. Annu Rev Genet 48, 187–214.

Kellis, M., Patterson, N., Endrizzi, M., Birren, B., and Lander, E.S. (2003). Sequencing and comparison of yeast species to identify genes and regulatory elements. Nature 423, 241–254.

Kleckner, N. (2006). Chiasma formation: chromatin/axis interplay and the role(s) of the synaptonemal complex. Chromosoma 115, 175–194.

Knop, M. (2006). Evolution of the hemiascomycete yeasts: on life styles and the importance of inbreeding. Bioessays 28, 696–708.

Kugou, K., Fukuda, T., Yamada, S., Ito, M., Sasanuma, H., Mori, S., Katou, Y., Itoh, T., Matsumoto, K., Shibata, T., et al. (2009). Rec8 guides canonical Spo11 distribution along yeast meiotic chromosomes. Mol Biol Cell 20, 3064–3076.

Kumar, R., Bourbon, H.M., and de Massy, B. (2010). Functional conservation of Mei4 for meiotic DNA double-strand break formation from yeasts to mice. Genes Dev 24, 1266–1280.

Kumar, R., Ghyselinck, N., Ishiguro, K., Watanabe, Y., Kouznetsova, A., Hoog, C., Strong, E., Schimenti, J., Daniel, K., Toth, A., et al. (2015). MEI4 - a central player in the regulation of meiotic DNA double-strand break formation in the mouse. J Cell Sci 128, 1800–1811.

Lacefield, S., and Ingolia, N. (2006). Signal transduction: external signals influence spore-number control. Curr Biol 16, R125–127.

Lam, I., and Keeney, S. (2015). Nonparadoxical evolutionary stability of the recombination initiation landscape in yeast. Science 350, 932–937.

Lange, J., Pan, J., Cole, F., Thelen, M.P., Jasin, M., and Keeney, S. (2011). ATM controls meiotic double-strand-break formation. Nature 479, 237–240.

Lange, J., Yamada, S., Tischfield, S.E., Pan, J., Kim, S., Zhu, X., Socci, N.D., Jasin, M., and Keeney, S. (2016). The landscape of mouse meiotic double-strand break formation, processing, and repair. Cell 167, 695– 708 e616.

Li, J., Hooker, G.W., and Roeder, G.S. (2006). Saccharomyces cerevisiae Mer2, Mei4 and Rec114 form a complex required for meiotic double-strand break formation. Genetics 173, 1969–1981.

Maleki, S., Neale, M.J., Arora, C., Henderson, K.A., and Keeney, S. (2007). Interactions between Mei4, Rec114, and other proteins required for meiotic DNA double-strand break formation in Saccharomyces cerevisiae. Chromosoma 116, 471–486.

Mancera, E., Bourgon, R., Brozzi, A., Huber, W., and Steinmetz, L.M. (2008). High-resolution mapping of meiotic crossovers and non-crossovers in yeast. Nature 454, 479–485.

Mimitou, E.P., Yamada, S., and Keeney, S. (2017). A global view of meiotic double-strand break end resection. Science 355, 40–45.

Miyoshi, T., Ito, M., Kugou, K., Yamada, S., Furuichi, M., Oda, A., Yamada, T., Hirota, K., Masai, H., and Ohta, K. (2012). A central coupler for 9 recombination initiation linking chromosome architecture to S phase checkpoint. Mol Cell 47, 722–733.

Mohibullah, N., and Keeney, S. (2017). Numerical and spatial patterning of yeast meiotic DNA breaks by TelGenome research 27, 278–288.

Murakami, H., Borde, V., Nicolas, A., and Keeney, S. (2009). Gel electrophoresis assays for analyzing DNA double-strand breaks in Saccharomyces cerevisiae at various spatial resolutions. Methods Mol Biol 557, 117–142.

Murakami, H., and Keeney, S. (2008). Regulating the formation of DNA double-strand breaks in meiosis. Genes Dev 22, 286–292.

Murakami, H., and Keeney, S. (2014). Temporospatial Coordination of Meiotic DNA Replication and Recombination via DDK Recruitment to Replisomes. Cell 158, 861–873.

Neale, M.J., and Keeney, S. (2006). Clarifying the mechanics of DNA strand exchange in meiotic recombination. Nature 442, 153–158.

Neale, M.J., and Keeney, S. (2009). End-labeling and analysis of Spo11oligonucleotide complexes in Saccharomyces cerevisiae. Methods Mol Biol 557, 183–195.

Padmore, R., Cao, L., and Kleckner, N. (1991). Temporal comparison of recombination and synaptonemal complex formation during meiosis in *S. cerevisiae*. Cell 66, 1239–1256.

Page, S.L., and Hawley, R.S. (2003). Chromosome choreography: the meiotic ballet. Science 301, 785–789.

Pan, J., Sasaki, M., Kniewel, R., Murakami, H., Blitzblau, H.G., Tischfield, S.E., Zhu, X., Neale, M.J., Jasin, M., Socci, N.D., et al. (2011). A hierarchical combination of factors shapes the genome-wide topography of yeast meiotic recombination initiation. Cell 144, 719–731.

Panizza, S., Mendoza, M.A., Berlinger, M., Huang, L., Nicolas, A., Shirahige, K., and Klein, F. (2011). Spo11-accessory proteins link double-strand break sites to the chromosome axis in early meiotic recombination. Cell 146, 372–383.

Raudsepp, T., Das, P.J., Avila, F., and Chowdhary, B.P. (2012). The pseudoautosomal region and sex chromosome aneuploidies in domestic species. Sex Dev 6, 72–83.

Robert, T., Nore, A., Brun, C., Maffre, C., Crimi, B., Bourbon, H.M., and de Massy, B. (2016). The TopoVIB-Like protein family is required for meiotic DNA double-strand break formation. Science 351, 943–949.

Rockmill, B., Voelkel-Meiman, K., and Roeder, G.S. (2006). Centromere-proximal crossovers are associated with precocious separation of sister chromatids during meiosis in Saccharomyces cerevisiae. Genetics 174, 1745–1754.

Sasanuma, H., Hirota, K., Fukuda, T., Kakusho, N., Kugou, K., Kawasaki, Y., Shibata, T., Masai, H., and Ohta, K. (2008). Cdc7-dependent phosphorylation of Mer2 facilitates initiation of yeast meiotic recombination. Genes Dev 22, 398–410.

Scannell, D.R., Zill, O.A., Rokas, A., Payen, C., Dunham, M.J., Eisen, M.B., Rine, J., Johnston, M., and Hittinger, C.T. (2011). The awesome power of yeast evolutionary genetics: New genome sequences and strain resources for the *Saccharomyces sensu stricto* genus. G3 (Bethesda) 1, 11–25.

Smagulova, F., Gregoretti, I.V., Brick, K., Khil, P., Camerini-Otero, R.D., and Petukhova, G.V. (2011). Genome-wide analysis reveals novel molecular features of mouse recombination hotspots. Nature 472, 375–378.

Stanzione, M., Baumann, M., Papanikos, F., Dereli, I., Lange, J., Ramlal, A., Trankner, D., Shibuya, H., de Massy, B., Watanabe, Y., et al. (2016). Meiotic DNA break formation requires the unsynapsed chromosome axis-binding protein IHO1 (CCDC36) in mice. Nat Cell Biol 18, 1208–1220.

Subramanian, V.V., Zhu, X., Markowitz, T.E., Vale Silva, L.A., San-Segundo, P., Hollingsworth, N.M., Keeney, S., and Hochwagen, A. (2018). Persistent DNA-break potential near telomeres increases initiation of meiotic recombination on short chromosomes. bioRxiv.

Sun, X., Huang, L., Markowitz, T.E., Blitzblau, H.G., Chen, D., Klein, F., and Hochwagen, A. (2015). Transcription dynamically patterns the meiotic chromosome-axis interface. eLife 4.

Taxis, C., Keller, P., Kavagiou, Z., Jensen, L.J., Colombelli, J., Bork, P., Stelzer, E.H., and Knop, M. (2005). Spore number control and breeding in *Saccharomyces cerevisiae*: a key role for a self-organizing system. J Cell Biol 171, 627–640.

Tesse, S., Bourbon, H.M., Debuchy, R., Budin, K., Dubois, E., Liangran, Z., Antoine, R., Piolot, T., Kleckner, N., Zickler, D., et al. (2017). Asy2/Mer2: an evolutionarily conserved mediator of meiotic recombination, pairing, and global chromosome compaction. Genes Dev 31, 1880– 1893.

Thacker, D., Mohibullah, N., Zhu, X., and Keeney, S. (2014). Homologue engagement controls meiotic DNA break number and distribution. Nature 510, 241–246.

Turney, D., de Los Santos, T., and Hollingsworth, N.M. (2004). Does chromosome size affect map distance and genetic interference in budding yeast? Genetics 168, 2421–2424.

Vincenten, N., Kuhl, L.M., Lam, I., Oke, A., Kerr, A.R., Hochwagen, A., Fung, J., Keeney, S., Vader, G., and Marston, A.L. (2015). The kinetochore prevents centromere-proximal crossover recombination during meiosis. Elife 4.

Wan, L., Niu, H., Futcher, B., Zhang, C., Shokat, K.M., Boulton, S.J., and Hollingsworth, N.M. (2008). Cdc28-Clb5 (CDK-S) and Cdc7-Dbf4 (DDK) collaborate to initiate meiotic recombination in yeast. Genes Dev 22, 386–397.

Wojtasz, L., Daniel, K., Roig, I., Bolcun-Filas, E., Xu, H., Boonsanay, V., Eckmann, C.R., Cooke, H.J., Jasin, M., Keeney, S., et al. (2009). Mouse HORMAD1 and HORMAD2, two conserved meiotic chromosomal proteins, are depleted from synapsed chromosome axes with the help of TRIP13 AAA-ATPase. PLoS Genet 5, e1000702.

Zhang, L., Kim, K.P., Kleckner, N.E., and Storlazzi, A. (2011). Meiotic double-strand breaks occur once per pair of (sister) chromatids and, via Mec1/ATR and Tel1/ATM, once per quartet of chromatids. Proc Natl Acad Sci U S A 108, 20036–20041.

Zhu, X., and Keeney, S. (2015). High-resolution global analysis of the influences of Bas1 and Ino4 transcription factors on meiotic DNA break distributions in *Saccharomyces cerevisiae*. Genetics 201, 525–542.

## References

Bao, Z., Xiao, H., Liang, J., Zhang, L., Xiong, X., Sun, N., Si, T., and Zhao, H. (2015). Homology-integrated CRISPR-Cas (HI-CRISPR) system for one-step multigene disruption in *Saccharomyces cerevisiae*. ACS Synth Biol 4, 585–594.

Reid, R.J., Sunjevaric, I., Keddache, M., and Rothstein, R. (2002). Efficient PCR-based gene disruption in Saccharomyces strains using intergenic primers. Yeast 19, 319–328.

